# Genomic evidence of a single domestication origin for modern cattle

**DOI:** 10.64898/2026.11.04.764331

**Authors:** John L. Miraszek, Alan Cooper, Bastien Llamas, Kefei Chen, Kieren J. Mitchell, Ayla van Loenen, Holly Heiniger, Yassine Souilmi, Graham Gower, Harly J. Rowan, Patrick J. Wrinn, Edgar A. Diaz Miranda, Simone Eliza Facioni Guimarães, Sergei Vasil’ev, Nikoli D. Ovodov, Jeremy F. Taylor, Robert D. Schnabel, Jared E. Decker

**Author notes:** Correspondence and requests for materials should be addressed to Jared Decker, regarding overall paper and population genomic analyses, Robert Schnabel regarding sequence alignment and variant calling, and Alan Cooper regarding ancient DNA sequencing. Curtin Biometry and Agricultural Data Analytics, School of Molecular and Life Sciences, Curtin University, Bentley, Perth, WA 6102, Australia.

## Abstract

Domesticated cattle occur as humpless taurine and humped zebu, historically explained as two independent domestications in the Fertile Crescent and Indus Valley. We question the validity of this model using whole-genome analyses of cattle and their wild relatives, including extinct aurochs. Zebu derives from admixture between domesticated taurine cattle and Indian aurochs and has elevated genomic diversity from an ancient introgression of a gaur-like or banteng-like population into Indian aurochs. We identify selective introgression in zebu from both 238 Indian aurochs-derived genes enriched for environmental adaptation, and 283 taurine-derived genes enriched for domestication loci. We suggest this process was shaped by polygenic selection. Modern cattle diversity traces from a single domestication event, followed by migration, wild introgression, and selection.

## Introduction

Cattle were among the first large mammals to be domesticated and were important to early human agrarian societies^1^. For the last 30 years it has been accepted that there were two independent domestications of cattle^2^, the first in the Fertile Crescent from the wild aurochs ancestor (*Bos primigenius*) and the second in the Indus Valley from Indian aurochs (*Bos primigenius namadicus*). The two extant biological types are commonly referred to as subspecies: *Bos taurus taurus* (Fig. 1b) and *Bos taurus indicus* (Fig. 1c), referred to as taurine and zebu, respectively. The subspecies are easily distinguished by morphological differences, including in zebu: a large hump on the neck, increased skin hanging from the neck and navel, large, pendulous ears, and increased size and density of sweat glands^3–5^ (Fig.1c). Further, there is a considerable difference in genome-wide nucleotide diversity (π) between the subspecies (zebu average one single nucleotide polymorphism (SNP) per 285 base pairs, taurines average just one SNP per 714 base pairs^6^). Moreover, the mitochondrial genomes of taurine and zebu are estimated to have diverged 200,000 to 1 million years ago, long before human domestication of animal and plant species^7^. These observations have been inferred to support the model of two independent cattle domestications. However, the conclusion that zebu were independently domesticated is heavily based on the mitochondrial data, and the evolutionary history of the zebu autosomal genome has not been resolved (Fig. 2A).

**Fig. 1.**
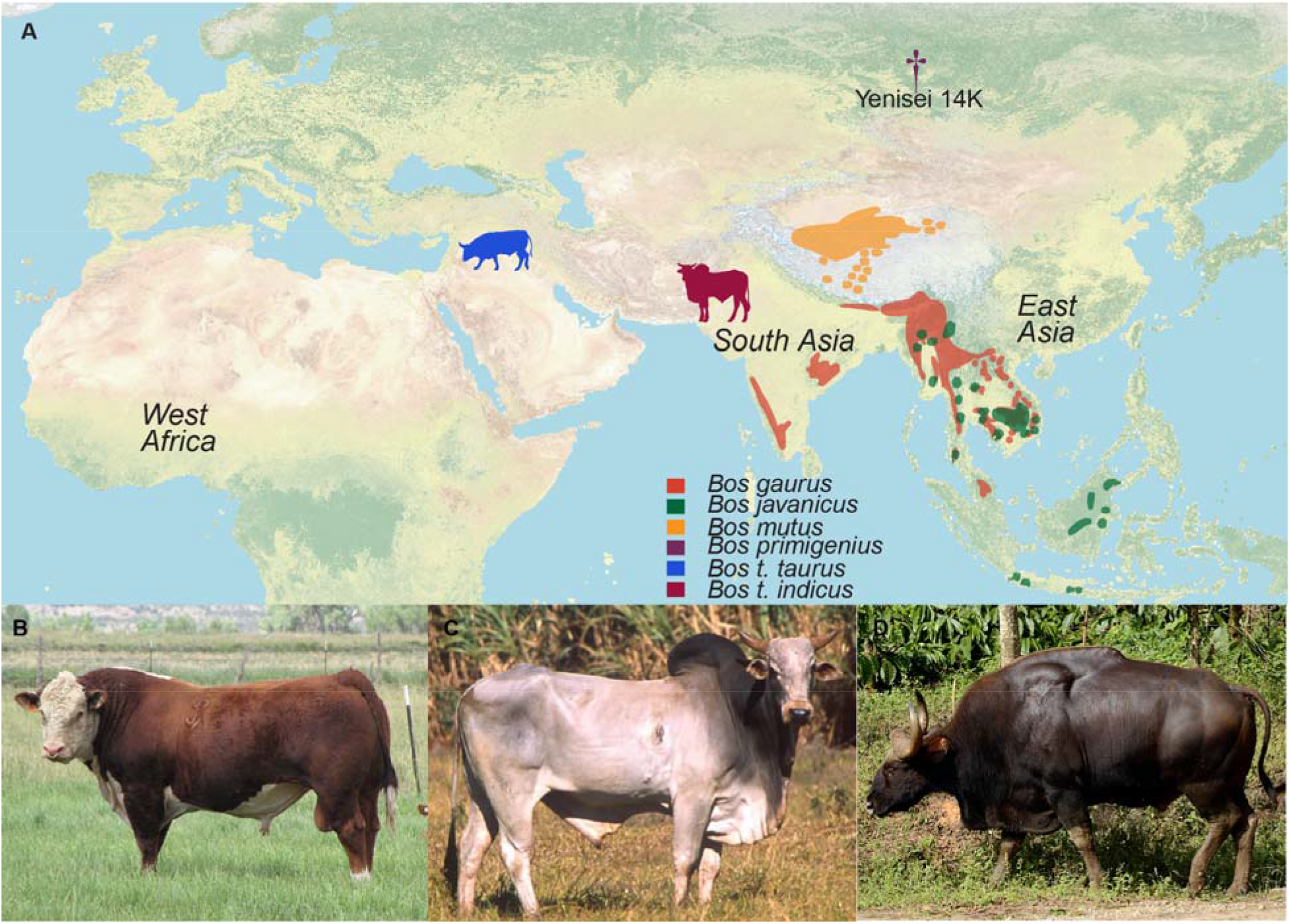
Geographic locations of wild and domesticated cattle. (**A)** Geographic distributions of modern *Bos gaurus* (gaur), *Bos javanicus* (banteng), and *Bos mutus* (wild yak), with the Fertile Crescent denoted by a blue taurine and the Indus River Valley marked by a magenta zebu. The location of the Yenisei 14K *Bos primigenius* aurochs sample sequenced in this research is denoted by a purple dagger. Other key locations of admixture are denoted with text labels. Geographic distribution data retrieved from the Map of Life^63,64^. Taurine silhouette by Mariana Ruiz Villarreal obtained via PhyloPic (Public Domain Mark 1.0). Zebu silhouette by T. Michael Keesey obtained via PhyloPic (CC0 1.0). Public domain map from Natural Earth, obtained using the rnaturalearth package. **(B)** Representative taurine bull, photo by Sue Decker. **(C)** Representative Zebu bull, photo by Scott Bauer, public domain via USDA Agriculture Research Service. **(D)** Representative *Bos gaurus* bull, photo by P. Jeganathan (CC BY-SA 4.0).

**Fig. 2.**
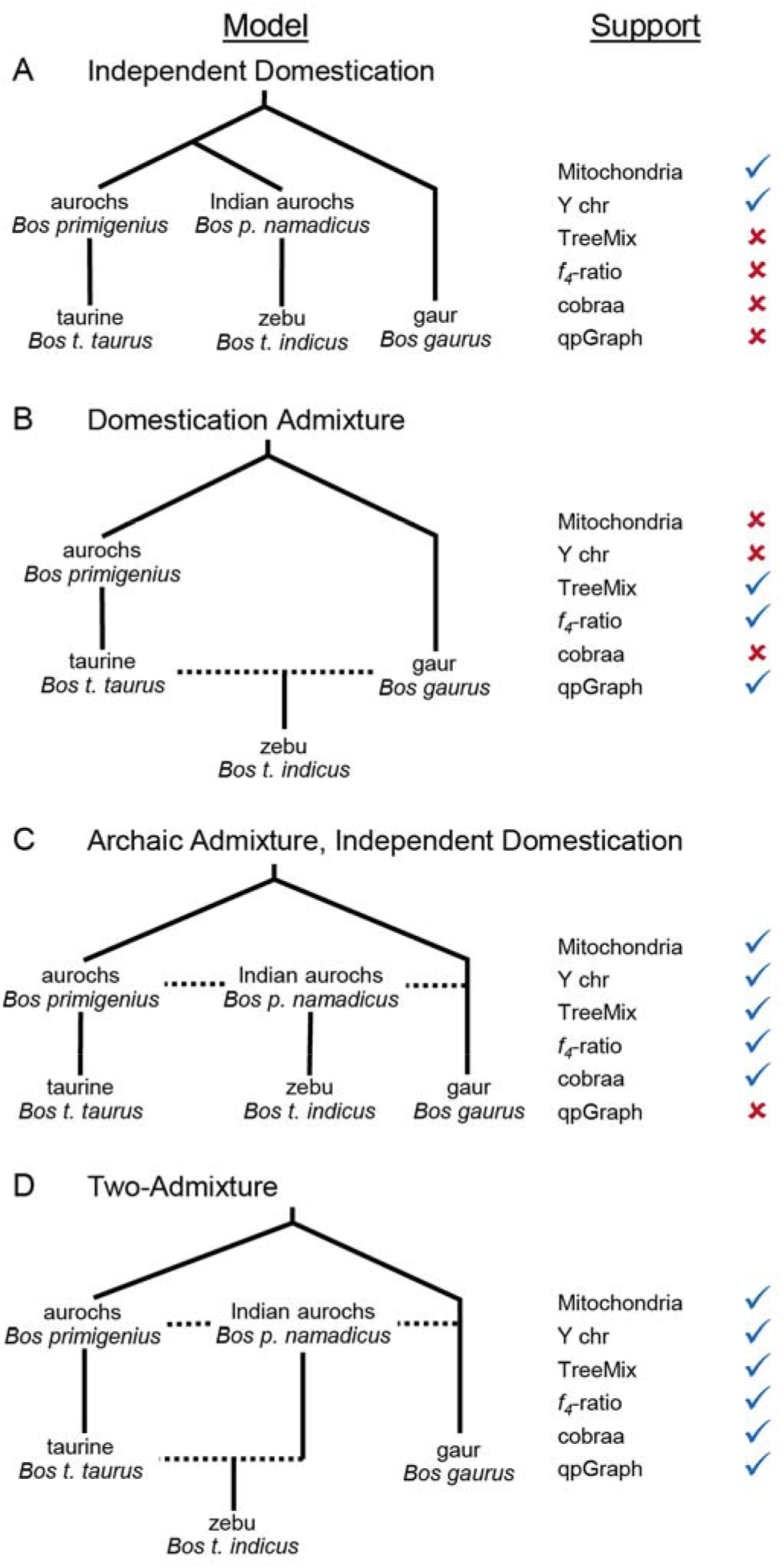
Competing models of zebu domestication, with supporting data and analyses designated. Check marks indicate congruent support; x marks indicate data and analyses that reject that model. Dashed lines represent admixture. **(A)** Independent domestication in the Fertile Crescent of taurine and Indus River Valley of zebu. **(B)** Domestication Admixture model in which admixture between taurine and gaur produce domesticated zebu. **(C)** Archaic admixture between aurochs and gaur to produce Indian aurochs. In this model, zebu result from independent domestication of admixed Indian aurochs. **(D)** Two-admixture model in which archaic admixture between aurochs and gaur produce Indian aurochs, followed by historical admixture between taurine and Indian aurochs to produce zebu.

The incorporation of population genomic principles into research focusing on the domestication of different species has resulted in a single domestication event becoming the default null hypothesis^8^. Strong evidence to the contrary is required to conclude that multiple independent domestication processes occurred within a species ^8^. This is particularly important because a history of admixture is common in the formation of differentiated domesticated populations, a process which mirrors the admixture of archaic^9–12^ and historical^13–15^ human populations. Post-domestication admixture between domesticates and their ancestral wild populations has repeatedly been identified, *e*.*g*., in pigs^16^ and chickens^17^, overturning earlier interpretations that multiple domestication events occurred in these species. Further, the ancestry from wild admixture donors can even replace the ancestry contributed by the initial domesticated populations^16^.

While animal domestication has become better understood, the genes and phenotypes selected during domestication remain poorly understood. Population genomic analyses have historically struggled to identify loci involved in polygenic selection, and this can lead to prioritizing the identification of selective sweeps over searching for polygenic adaptation^18^. Comparisons to wild relatives in plant domestication scenarios have revealed large-effect loci related to key domestication phenotypes (*i*.*e*., loss of seed shattering) which serve as the most straightforward examples of ‘domestication genes’^19^ . In the case of animal populations, the pathways essential to fitness in a domesticated niche are more complex and polygenic, especially for behavioral, metabolism, and health phenotypes. Specifically, for adaptation during the earliest phases of domestication, there are few examples of domestication selective sweeps in animals^20^. Previously proposed animal domestication loci are variable in sequenced ancient genomes and reflect modern selection pressure^20^. We approach this issue by leveraging the power of methods that identify adaptive introgression, analyzing patterns of allele frequencies rather than relying on selective sweep methods. Further, we assess selective introgression from both the wild bovid and domesticated cattle donors during early domestication, as farmers created domesticated populations across South Asia, Africa, and East Asia.

## Results

### Global genetic diversity in the genus *Bos*

To assess the population history of cattle domestication, we utilized a cohort of genome sequences for 5,606 animals^21^ (Table S1). The genome sequences were all identically processed for variant calling (Methods) and from this cohort, a subset of 1,804 animals was chosen for further analysis to reduce the over-representation of commonly sequenced populations. The analyzed data represented 5 species groups: bison (*Bison bison*), wild yak (*Bos mutus*) and their domesticated counterpart yak (*Bos grunniens*), gaur (*Bos gaurus*) and their domesticated counterpart gayal (*Bos frontalis*), banteng (*Bos javanicus*), aurochs (*Bos primigenius*) and their domesticated counterparts taurine and zebu cattle (Fig. 1a-d). Aurochs ancestry was represented by an 18X coverage genome sequence for a specimen recovered from the Yenisei River in Russia, which was dated to 14,000 years old (referred to as Yenisei 14K). The analyzed animals represent over 230 populations and breeds. All extant members of the genus *Bos* produce fertile females when crossed^22^, and there are many documented cases of interspecific gene flow in this clade^23^. Consequently, sampling the breadth of global diversity better represents the totality of genetic ancestry for these populations than does geographically limited or species-based cohorts. We included diversity from multiple species in variant quality score recalibration to avoid reference bias. After quality control, including only biallelic SNPs, and linkage disequilibrium pruning (see Methods), 38,097,187 SNPs were retained for analyses.

### Origin of domesticated zebu cattle

We performed exploratory^24^ principal component^25^ and ancestry assignment^26^ analyses for subsets of the 1,804 sequenced *Bos* animals (Figs. S1-S3). These confirmed the identity of the samples, and results were congruent with previous results using SNP array genotypes^6,27,28^. TreeMix^29^ was used to investigate phylogenetic relationships and admixture events across alternative data subsets (Fig. S4) and consistently revealed strong support for admixture between the gaur-gayal clade and the base of the clade of Asian zebu populations (ancestry proportion = 0.144, p-value < 2.2e-308, Fig 2B). In fact, this gaur to zebu gene flow was the first migration edge added to various subsets of the global *Bos* phylogenetic network in all but one model (Fig. S4), where it was second to banteng introgression into East Asian zebu.

To further investigate the source of the admixture, we used *f*_*3*_ statistics, *f*_*4*_-ratios, and admixture graphs^30^ to test how admixture between taurine and a gaur-like population could have formed the progenitor of zebu. There were no significant *f*_*3*_ statistics where any of the Indian subcontinent zebu breeds (Sahiwal, Gir, Hariana or their derivatives Nelore and Brahman) resulted from admixture between gaur or their sister species banteng and a taurine population (Table S2). Since the *f*_*3*_ signal erodes over time due to genetic drift^30^, the lack of significant *f*_*3*_ results suggests that the gaur-like admixture with the aurochs’ ancestors of zebu was an archaic event. The *f*_*4*_-ratio test is more powerful for detecting ancient admixture^30^, and we found that when domesticated taurine cattle formed one ancestral population, *f*_*4*_-ratios were significant and estimated that modern zebu had received 40% (Z-score = 193.46) to 51% (Z-score = 147.866) of their ancestry from a gaur-like or banteng-like population (Table S3).

To further resolve the admixture history between cattle and gaur in the zebu domestication process we explored numerous admixture graph topologies. We first compared two competing models of zebu ancestry. The Independent Domestication Model (Fig. 2A) was a bifurcating phylogeny representing separate domestications with an old divergence between zebu and taurine cattle (predating the split between the 14,000-year-old Yenisei aurochs sample and taurine, Fig. S5a). The Domestication Admixture (Fig. 2B) graph modeled a hybrid origin for zebu as a mixture of taurine and gaur-like ancestry (Fig. S5b) and had a significantly better fit (empirical p-value < 0.0002, Table S4), indicating zebu clearly have an interspecific admixed origin.

We next utilized cobraa^31^, an extension of Pairwise Sequentially Markovian Coalescent (PSMC), which applies a coalescence-based hidden Markov model to analyze individual genomes and infer effective population sizes over time and the timing and proportions of admixture events.

We used the three highest sequence coverage zebu samples from South Asia to assess the consistency of results. The cobraa analyses identified several scenarios of deep population structure (*i*.*e*., included admixture) which were superior to panmictic analyses (no admixture) (Fig. S6). The best-fit models estimated that the two populations contributing to the admixture had divergence times of 4 to 8 million years. However, the timing of the admixture itself (estimated between 51,933 and 209,865 generations ago or 259,667 to 1,049,326 years ago, Fig. S6-S9) was much older than the domestication of cattle. This indicates that the Indian aurochs itself was the product of archaic admixture between two ancestral populations (Fig. 2C). The estimated archaic admixture was estimated to be 22%, 28%, and 47% (Fig. S6), which agrees with the TreeMix graph estimates of gaur ancestry and is consistent with the mitochondrial divergence between taurine and zebu. Compared to *f*_*4*_-ratios with domesticated cattle as the donor, the *f*_*4*_-ratio estimates in which aurochs were the donor population estimate smaller contributions (34-39%) from the gaur-like or banteng-like ancestor (Table S3), which are more similar to estimates from cobraa and TreeMix, further supporting archaic admixture.

To explore the potential of multiple admixture events, we tested two competing admixture scenarios. The Archaic Admixture Independent Domestication model (Fig. 2C) contains archaic admixture between gaur and aurochs to produce Indian aurochs, which was subsequently independently domesticated to produce zebu (Fig. S5c). Alternatively, a Two-Admixture Model (Fig. 2D) features admixture between gaur and aurochs to produce Indian aurochs, followed by a second admixture event between Indian aurochs and taurine to form the domesticated zebu population (Fig. S5d). Despite the increased network complexity, the Two-Admixture model provided a significantly better fit to the data (empirical p-value < 0.0002, Table S4). It is also congruent with the Y chromosome data (Fig. S10), and previously published analyses of Y chromosome^32^ and mtDNA haplogroups^33–36^, where zebu haplotypes are notably divergent but are more similar to taurine and aurochs than to other *Bos* species (Figs. 2, 3).

**Fig. 3.**
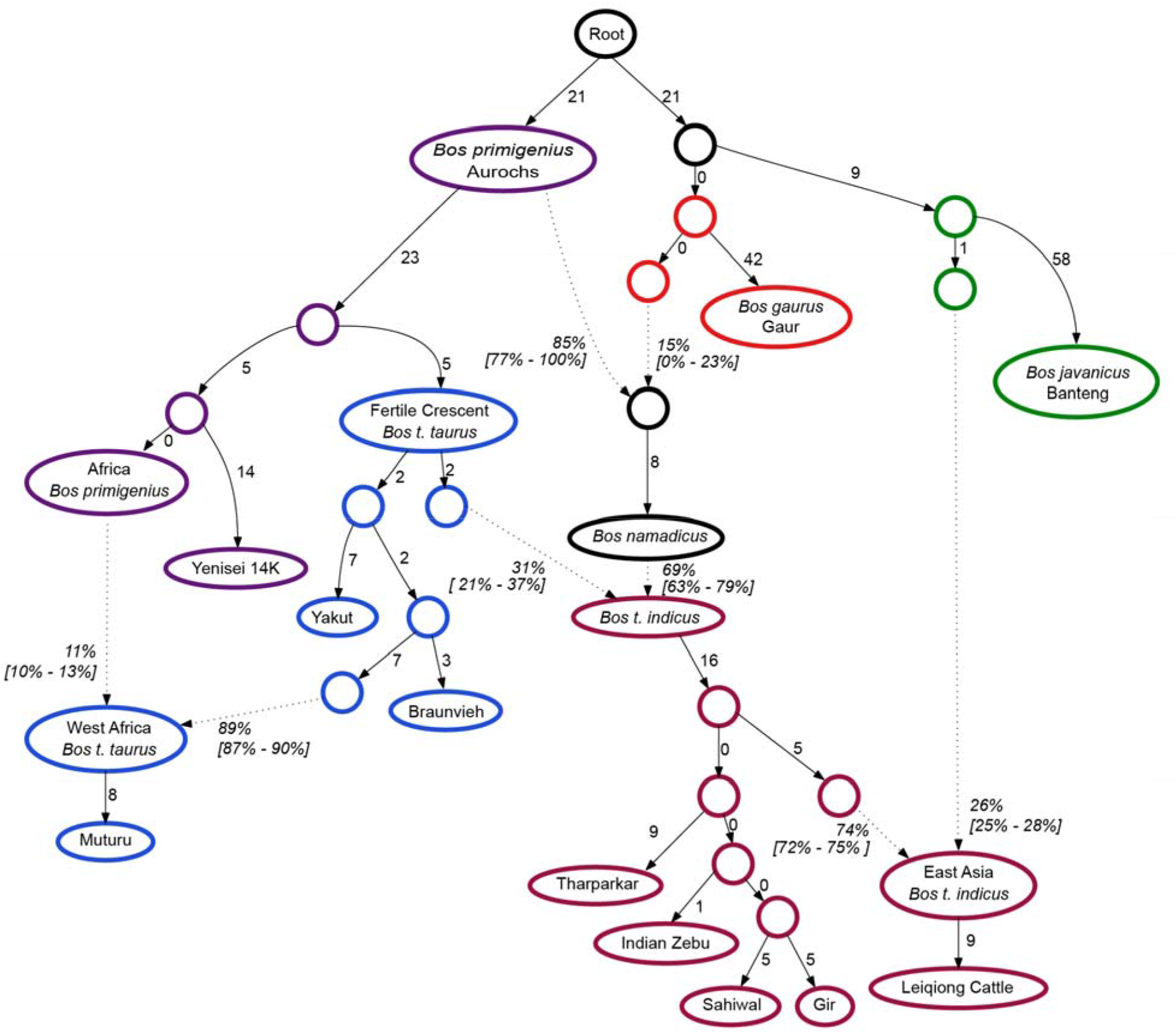
Admixture graph of South Asia, Africa, and East Asia cattle populations. Admixture graph of two waves of admixture for zebu, aurochs introgression into taurine in West Africa, and *Bos javanicus* (banteng) introgression into zebu in East Asia. Values in square brackets reflect 95% confidence intervals. Graph fit statistics are in Table S4.

Numerous qpGraphs topologies were tested, moving the location of the donor nodes in the network (Fig. S11 – S21). Admixture coming directly from the edge leading to the Yenisei 14K sample and a wild *Bos* ancestor estimated less than 50% of ancestry of Indian aurochs coming from aurochs (Fig. S11), which is not congruent with Y chromosome (Fig. S10) or mtDNA data ^33–36^. Admixture between the Yenisei branch and ancestral gaur produced reasonable estimates of aurochs ancestry for the first admixture event, but the proportion of taurine ancestry was much higher than other models (Fig. S12). The graph plotted in Fig S13 is also not probable considering the Y chromosome and mtDNA data. Several models produced ancestry estimates of 0% from wild *Bos* (Fig. S14-S16), which disagree with cobraa results. Admixture graphs plotted in Figs S12, S17, and S18 had extremely poor fit scores. The graphs are plotted in Figs. S19 to S21 have the strongest agreement with our other results, with Fig S19 having the best score of graphs inferring both archaic and ancient admixture.

### Adaptive introgression in zebu

To identify selected adaptive loci introgressed into modern zebu cattle, we calculated *f*_*dM*_ statistics^37,38^ which measure the extent to which allele frequencies at a given locus are tree-like versus admixed-like (an extension of Patterson’s D used for genome-wide scans). Because this method allows for the explicit specification of donor populations, we calculated *f*_*dM*_ twice; once with a wild donor (Fig. 4a), and once with a domesticated taurine donor (Fig. 4c). Previous searches for loci selected during domestication have focused on methods which identify selective sweeps. However, hard sweeps are now thought to be the exception rather than the rule, while in contrast polygenic selection can create rapid changes in the means of selected traits, via subtle shifts in allele frequencies at thousands of loci^39,40^. Rapid trait changes would have been necessary to successfully move animals from a wild to a domesticated niche. Allele frequency correlations manifesting as large introgression statistics (such as *f*_*dM*_) provide an opportunity to identify loci exposed to polygenic selection due to forces like domestication. From the genome-wide scans, we identified functional annotations for the genes within or overlapping SNP (∼3,250 bp) windows with the 99.5^th^ highest percentile of *f*_*dM*_ scores (Figs. 4b and 4d). Positive *f*_*dM*_ statistics reflect strong signals of admixture from the donor population, while negative *f*_*dM*_ values indicate shared variation between the sister and donor populations^37^. *A priori*, we might expect that enriched annotations and associations of genes within these windows would be related to traits specific to the niche of each donor (*e*.*g*., local environmental adaptation for Indian aurochs/gaur-introgressed alleles, and domestication associated adaptation for taurine-introgressed alleles). Gaur samples were used as a surrogate for Indian aurochs ancestry, based on TreeMix results. We identified 238 genes (116 with gene symbols) in selected regions with an Indian aurochs/gaur origin and 283 genes (147 with gene symbols) introgressed from domesticated taurine. For Indian aurochs introgression, for genes without gene symbols, 22 were lncRNA, 2 were miRNA, 41 were protein coding, 3 were pseudogenes, and 2 were TR V genes. For domesticated taurine introgression, for genes without gene symbols, 31 were lncRNA, 1 was a miRNA, 49 were protein coding, 3 were pseudogenes, and 3 were TR V genes.

**Fig. 4.**
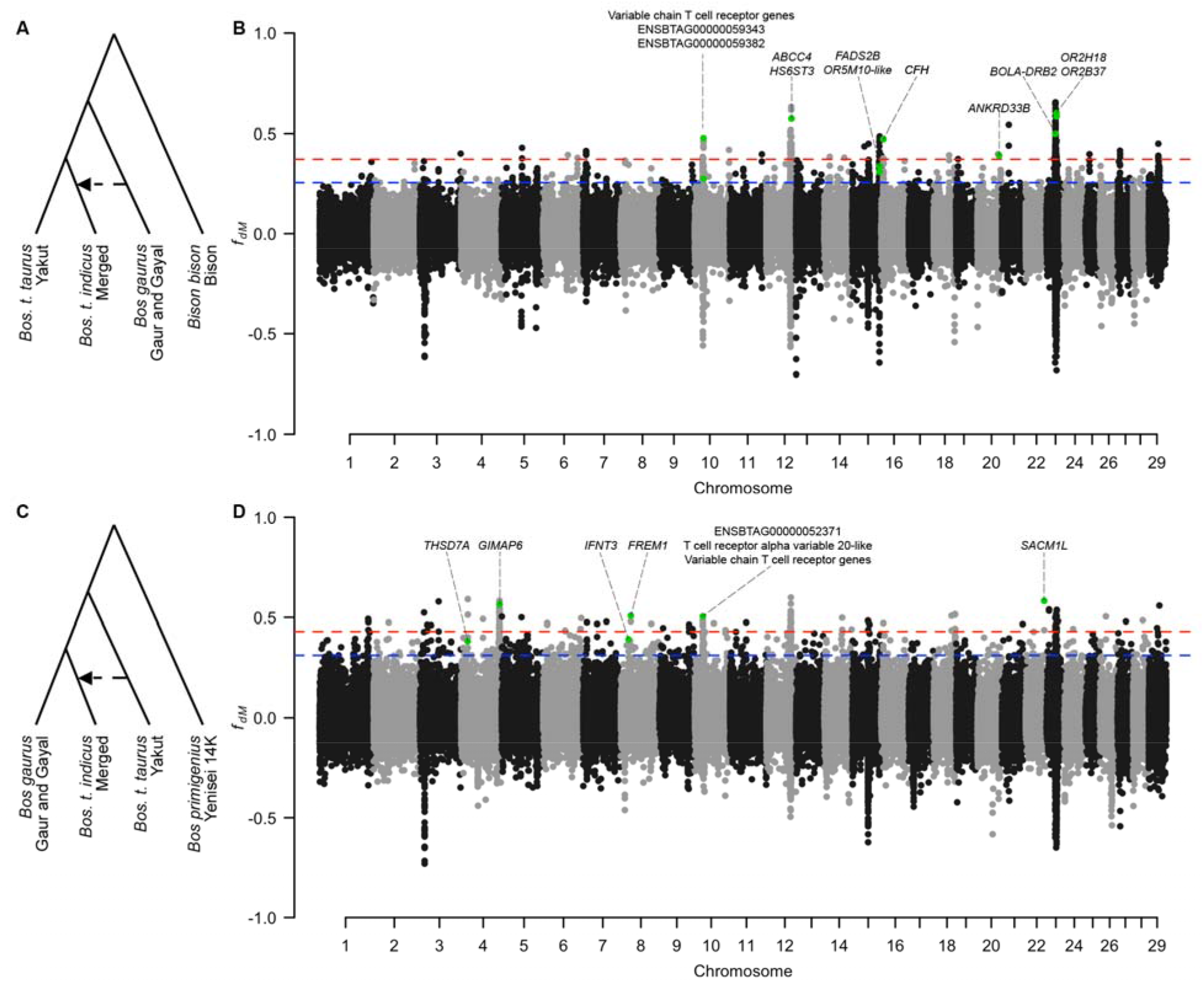
Genome-wide scans of adaptive introgression in zebu from South Asia. **(A)** Phylogeny and populations used in *f*_*dM*_ analysis of wild introgression into zebu. **(B)** Manhattan plot of *f*_*dM*_ values for wild introgression into zebu. (**C)** Phylogeny and populations used in the *f*_*dM*_ analysis of domesticated cattle introgression into zebu. (**D)** Manhattan plot of *f*_*dM*_ values for domesticated introgression into zebu. Dashed blue line represents the cutoff for the 99.5^th^ percentile and the dashed red line indicates the cutoff for the 99.9^th^ percentile. Loci above the 99.5^th^ percentile were used to investigate enrichment. Loci containing discussed candidate genes are highlighted in green.

Strong signals of Indian aurochs/gaur introgression were observed on 7 chromosomes (Fig. 4b, supplementary text S1), and the 116 genes within introgressed loci were enriched for functions related to immune; metabolic and endocrine; cognitive; cardiovascular and hematological; respiratory; liver and gastrointestinal; and musculoskeletal systems (Table S5).

The functions of loci selectively-introgressed from taurine ancestry included behavioral and cognitive; immune; metabolic; eye; endocrine; birth and developmental; and growth and morphological traits (Table S6, supplementary text S1). These genes included transcription factor targets such as *SMAD4* (immune regulation and embryonic development), *TCF4* (brain development), *STAT3* (immune responses, cell growth, and metabolism), *WT1* (kidney, gonad, and heart development), and *ESR1* (female reproduction).

Although there were many gene annotation terms common to the introgressed gene lists from both Indian aurochs/gaur and taurine, only 24 of the 521 genes were shared. In some instances, shared genes and terms likely reflect balancing selection for adaptation to environmental challenges (*e*.*g*., immunity) for which increased heterozygosity and sequence diversity are advantageous. However, the shared enriched terms, especially those related to behavior, could reflect selection in opposing directions at different evolutionary time points. Zebu cattle are more aggressive and excitable^41^ than taurine cattle due to a stronger antipredator response^5,42^. As a wild prey animal, Indian aurochs would have derived a fitness advantage from increased predator avoidance behavior. The docility of taurines was undoubtedly strongly selected by humans following domestication, but any increase in frequency of docility alleles would have been reduced in the initial zebu due to admixture with wild Indian aurochs. Thus, the behavioral differences between modern taurine and zebu may reflect their different histories of admixture and post-domestication selection intensity.

Loci introgressed from the Indian aurochs/gaur ancestor were significantly enriched for gene expression in the cerebral cortex in Mouse ENCODE and the Human Protein Atlas data, and in the salivary gland, breast, and ovary in the human GTEx data. Loci introgressed from taurine were enriched for gene expression in the cerebral cortex in the Human Protein Atlas, fallopian tube, thyroid, brain, cervix uteri, and salivary gland in GTEx, and cortex, E14.5 brain, olfactory bulb, and cerebellum in Mouse ENCODE. These enrichments highlight the importance of the brain, and thus likely various behaviors in both admixture events. Further, the thyroid and female reproductive organs are important to domestication. Wild aurochs were seasonal breeders, whereas modern domesticated cattle can breed year-round^43^. Ablation of the thyroid gland results in the loss of seasonal breeding^44^. The enrichment of selected genes expressed in these tissues highlights the importance of reproduction in the transition from wild to domesticated niches^45,46^.

From our sequence variant calls, we identified a small number of missense mutations within the genes in these introgressed regions (supplementary text S2, Table S7), providing further evidence for the limited importance of selective sweeps during domestication. Of the 425 protein coding genes identified in the adaptive introgression analysis, only 6 had protein coding changes. and had the correct allele frequency concordance matching the history of admixture from the donor into zebu (Table S6). Of these, 2 were in olfactory receptor genes, a common target of adaptation and domestication^47,48^. Sweeps frequently act on large effect protein coding changes, whereas polygenic selection acts on smaller effect variants. These results are congruent with selection for local adaptation and domestication in the evolutionary history of modern zebu cattle that was polygenic in nature and did not rely on hard sweeps on protein coding changes.

### Wild introgression in Africa and East Asia

To examine the extent of adaptive introgression from wild relatives we used the admixture graph model to test additional previously reported admixture events in Africa^27,49^ and East Asia^27,50–52^ as early domesticated cattle were moved around the world by human herders (Fig. 3). Due to the genetic divergence between taurine cattle in Africa from those in Europe and Asia, it has been suggested that there may have been a third domestication of cattle in Africa^53,54^. Using our high-coverage ancient aurochs sample, Yenisei 14K, we find that the ancestry of African taurine cattle includes significant admixture from African aurochs (p-value < 0.0002, Fig. 3, Fig S22a, supplementary text S3). This explains the divergence of African taurine cattle from European and Asian domesticated cattle and refutes suggestions of an independent domestication of cattle in Africa.

In agreement with previous reports^27,50,51,55^, all of the analyses of East Asian cattle (TreeMix, admixture graphs, and *f*-statistics) detected an admixture history from banteng (supplementary text S4). Indeed, the banteng introgression into East Asian cattle was the single most impactful admixture event influencing the model fit in our world-wide model of cattle ancestry (p-value < 0.0002, Fig. 3, Fig. S22b). This is likely due to the recent nature of this admixture and the large divergence between the donor populations.

Known admixture between aurochs and European cattle was not included in reported models, as this requires additional degrees of freedom be utilized in our admixture graphs. However, this results in larger scores for our admixture models.

### Selection on loci following admixture identifies specific local adaptation and consistent selection for domestication

We compared the selectively-introgressed genes identified by *f*_*dM*_ analyses across the three admixture events in South Asia, Africa, and East Asia (Fig. S23, Fig. S24, supplementary text S1). For the 722 genes identified as originating from a wild donor (Indian aurochs, African aurochs, or banteng), none were common across all three admixture events and only 46 were identified in two of the admixture events. Just three gene annotation and association terms were significantly enriched across all three analyses (Table S8). The selected genes likely reflect selection for adaptation to local environments and are unique to their specific context. Enriched terms across the three wild-introgression analyses were typically related to immune functions, cardiovascular traits, body size and neurological traits. Examples include loci linked to critical traits such as heat tolerance in zebu and disease resistance in African taurine breeds (supplementary text S1).

For genes selectively-introgressed from a domesticated donor, one gene was common across all three events, 48 genes were shared by two events, and 980 were unique to a single event. In contrast to wild donors, there were 74 annotation terms that were significantly enriched across all three admixture events (Table S9, Fisher’s combined probability test p-values < 1e-04). Twenty-five of these terms were transcription factor targets, including *TCF4* which is involved in nervous system development^56^ and neurodevelopmental disorders^57^. Of the remaining 49 annotation terms, 17 are involved in behavioral traits. The convergence of phenotypes in different populations when selection acts on different sets of variable loci within a given pathway (genetic redundancy) is a defining feature of polygenic adaptation^40,58^. However, selective sweeps, especially those earliest in domestication, should be shared across breeds. When we analyzed adaptive introgression sweeps in South Asia zebu breeds, we found no sweeps that were shared across breeds. Rather, all three populations adapted to the niche of domestication following admixture via selection on different sets of variable genes within common pathways, a hallmark of polygenic selection.

## Discussion

Since the model of independent zebu domestication was first proposed it has rarely been challenged, and large-scale analyses of ancient aurochs genomes^35,59^ have not fully explained the diversity of zebu cattle. However, multiple lines of evidence from our data (TreeMix, *f*_*4*_-ratios, cobraa structured coalescent, and qpGraphs analyses) all refute a completely independent second domestication of zebu cattle^2,60^, and suggest that introgression of domestication-related alleles from previously domesticated taurines facilitated the formation of domesticated zebu cattle. Much like swine in Europe^16^, there was likely a nearly complete turnover from Fertile Crescent domesticated cattle mitochondrial haplotypes to Indian aurochs mitochondrial haplotypes in Indus Valley zebu cattle. However, to definitively conclude whether or not zebu were independently domesticated requires accurate dating of admixture between taurine and Indian aurochs population or ancient DNA from the time period of zebu domestication.

The domestication of cattle from large, wild progenitors was a prolonged, hazardous process, characterized by repeated hybridization with wild bovids. Cattle are likely similar to most domesticated animal species, arising from a single domestication process^8^ but with frequent gene flow from wild populations^16,17,23,61,62^. Multiple rounds of archaic and historic admixture within the *Bos* genus not only obscured the relationship between taurine and zebu lineages, but our results also explain the elevated nucleotide diversity^6^ observed in zebu cattle. The larger nucleotide divergence and diversity in Indian aurochs relative to Fertile Crescent aurochs has not to this point been understood, but our results account for this through an ancient admixture with another *Bos* species (similar to gaur or banteng). As Neolithic herders dispersed domesticated cattle out of the Fertile Crescent, cattle were exposed to novel climatic and pathogenic pressures. Introgression from locally adapted wild *Bos* populations provided an expeditious route to environmental adaptation. Our *f*_*dM*_ analyses suggest that the introgression of these adaptive alleles provided a rapid mechanism for domesticated cattle to acquire environmental resilience. Further, for domestication loci, the genetic signals do not show shared selective sweeps and support a more prevalent role of polygenic selection shifting allele frequencies genome wide.

Genes directly exposed to selection during the earliest phases of the domestication process are poorly understood. By leveraging genomic methods designed to detect adaptive introgression, it is possible to identify the genes and functions contributed by different wild and domestic ancestors and overcome historical biases that favor selective sweeps over polygenic selection. Because this framework infers adaptive introgression from historical admixture, it offers a generalizable framework for pinpointing genes involved in domestication and local adaptation, thus informing genomic selection and gene editing aimed at climate-resilient agricultural production.

## Acknowledgments

J.F.T. was supported by grants 2015-67015-23183 and 2017-67015-26760 from the National Institute of Food and Agriculture of the United States Department of Agriculture and 1R01HD084353 from the National Institute of Child Health and Human Development. J.E.D was supported by Agriculture and Food Research Initiative Competitive Grants no. 2016-68004-24827, 2021-67021-33448, and 2025-67015-44928 and through the Hatch capacity funding MO-HAAS0027 from the USDA National Institute of Food and Agriculture. R.D.S. was supported by Agriculture and Food Research Initiative Competitive Grants no. 2020-67015-31675, 2020-67015-30829, 2020-67015-31132, 2021-67021-34149, 2023-67015-39261 and through the Hatch capacity funding MO-HAAS0001. Fazendas do BASA and Mr. Evandro Guimarães for funding the genomic sequencing of the Gir animals. CAPES Process 88887.577470/2020-00 for funding E.A.D.M. scholarship. We appreciate Julien Soubrier for assistance while sequencing the Yenisei 14K sample.

## Author contributions

J.E.D. conceived the project, and together with J.L.M. carried it out. B.L., K.C., K.J.M., A.v.L., H.H., Y.S., G.G., A.C. performed laboratory work and analysis of the Yenisei 14K sample. N.D.O. (deceased) discovered the Yenisei 14K fossil, assisted in identification and curation of the sample. P.J.W. sampling. S.V. sampling and curation. R.D.S. performed data retrieval, genome alignment and variant calling. H.J.R. performed preliminary analyses of sequence data and population genomic analyses. E.A.D.M. and S.E.F.G. provided sequence data for Gir samples. J.E.D., J.F.T. and A.C. provided funding for sequencing of the Yenisei 14K sample. J.F.T. provided funding for modern cattle sequencing and advised on data analysis and interpretation. J.E.D. and J.L.M. wrote the original draft of the manuscript with J.F.T. reviewing and editing drafts. J.L.M and J.E.D. created visualizations of the data.

## Data Availability

DNA sequencing data are available via the NCBI Sequence Read Archive (SRA).

## Code Availability

This study used publicly available software, referenced throughout the main text and methods.

Reprints and permissions information is available at www.nature.com/reprints.

## Supplementary Materials

Materials and Methods

Supplementary Text

Figs. S1 to S24

Tables S1 to S9

