## Supplementary Information 1 for "Genomic evidence of a single domestication origin for modern cattle"

**The PDF file includes:**

Materials and Methods
Supplementary Text
References
Figs. S1 to S10

**Other Supplementary Information for this manuscript include the following:**

Tables S1 to S9

### Materials and Methods

Ancient Aurochs Genome

A tibia specimen was collected 15 km north of Krasnoyarsk, Siberia, Russia, along the Yenisei River. It was identified as *Bos primigenius* based on morphology. Radiocarbon dating was performed by INSTAAR - Laboratory for AMS Radiocarbon Preparation and Research (University of Colorado at Boulder). Collagen was isolated and purified using ion exchange chromatography. Graphite prepared from the purified amino acids fraction was dated using mass spectrometry to 12,300 ± 40 radiocarbon years BP ($\partial^{13}C_{wrt\ PDB}$: -17.7‰; AMS result number CURL-10282). Date calibration was performed using the IntCal 20 curve in OxCal 4.4.4[1]: mean ± σ: 14,302 ± 195 calBP; median: 14,237 calBP; 83.5% probability: 14,1440 – 14,083 calBP; 12.0% probability: 14,805 – 14,710 calBP.

All ancient DNA laboratory work was performed in purpose-built facilities at the University of Adelaide's Australian Centre for Ancient DNA (ACAD). To reduce contamination with exogenous DNA, sample surfaces were irradiated with UV for 15 min, then abraded using a Dremel™ rotative tool fitted with a carborundum cutting disc. Ancient DNA was isolated and purified from 220 mg of bone fragments as previously described[2]. A double-stranded Illumina sequencing library was built from 25 µL of DNA extract as previously described, using a partial uracil-DNA-glycosylase and endonuclease VIII (USER; New England Biolabs) treatment[3]. Library amplification was performed using AmpliTaq Gold DNA Polymerase (ThermoFisher Scientific). The mitochondrial DNA fraction was enriched using the Mybaits v.2 protocol (Daicel Arbor Biosciences) and sequenced on an Illumina MiSeq (Australian Genome Research Facility). Both short fragment length and presence of 5' C-to-T and 3' G-to-A substitutions were indicators of authentic ancient DNA. Upon molecular validation of the morphological assignment to *Bos primigenius*, another double-stranded library was prepared as above—this time with full USER treatment, and amplified in eight 25-µL replicate PCR reactions using Phusion™ high-fidelity DNA polymerase (New England Biolabs) and 6 cycles of amplification (the number of PCR cycles was estimated based on rtPCR, using the same polymerase). The library was shotgun sequenced across multiple flow cells on Illumina NextSeq 500 (Australian Genome Research Facility) and HiSeq 2500 (Queensland Brain Institute) platforms.

Whole genome data acquisition

This project used whole genome resequencing data from the previously described UMAGv1 bovine cohort[4]. Briefly, publicly available genomes with unique BioSample IDs were identified as belonging to domesticated cattle or wild bovid species based on their taxon IDs and were collected from NCBI's SRA database. To account for the heterogeneity of sample origin, sequencing platform and year of upload, several quality control measures were taken as described below.

Variant calling

The raw sequence reads were aligned to the ARS-UCD1.2 *Bos taurus* reference genome assembly[5] with the addition of the Btau5.0.1 Y chromosome assembly, using either BWA-MEM (v0.7.17)[6] or BWA-MEM2 (2.2.1)[7] as previously described[4]. BQSR was performed with GATK BaseRecalibrator[8] using an updated known variant set from the 1000 Bulls Project[4] with adjusted parameters to include variants from wild *Bos* species without over-correction. A variant call set of SNPs and INDELs was generated using GATK[9]. As in that analysis, VQSR parameters were fine-tuned using bovine trios, including variants from bison, banteng, gaur, and yaks. Only variants that passed tranche level 99.7 were retained.

Metadata compilation

The raw metadata for each sample were collected using the Biopython package Bio.Entrez[10] to access the NCBI API. Metadata were gathered from each sample's listing on SRA, BioSample, or BioProject. Additional information was manually curated from published supplementary tables, where available. Samples were assigned a population/breed label based on all collected metadata, with normalization of nomenclature for factors such as spelling, capitalization, hyphen usage, abbreviation, and language (unless populations had distinct geographic data to suggest isolation [*e.g.*, Brown Suisse (Germany) / Braunvieh (USA) / Brown Swiss (Switzerland)]).

Down sampling of individuals per population/breed

A subsample of the initial cohort was extracted that included only the 40 highest coverage genomes for those domesticated populations with more than 40 samples to avoid over-sampling of popular breeds. Samples with less than 3X coverage were excluded. The resulting subsample of 1,804 whole genomes included 1,523 individuals from domesticated cattle breeds, 279 individuals from wild *Bos* species (banteng, bison, gaur, yak) or domesticated derivatives of yak and gaur populations, and 2 ancient aurochs genomes, and these data were used for population genetic analyses.

Variant down sampling and pruning

Biallelic SNP data for the 1,804 genomes were pruned for linkage disequilibrium using bcftools-1.14 with one SNP of each pair with $r^2 > 0.8$ being removed, resulting in 38,097,187 pruned and filtered biallelic autosomal SNPs (henceforth referred to as the 38M SNP set).

The pruned 38M SNP set was randomly down sampled to 100K, 150K, and 200K SNP sets by taking every *n*th SNP in a linearized autosomal genome, where *n* was 38M/(100K, 150K, or 200K) rounded down to the nearest integer.

Principal component analyses and ADMIXTURE

We performed various principal component analyses for quality control. For these analyses, the 38M SNP set was input to Plink-1.9b7[11,12] for the 1,804 global cohort and taxa specific sub-cohorts[4]. The 100K, 150K and 200K SNP sets were analyzed in ADMIXTURE[13] with K ranging from 2-10 for the 1,804 global cohort. Increasing the number of SNPs analyzed in ADMIXTURE did not produce meaningful differences in the results, and consequently, the 100K SNP set was used for all further exploratory analyses. Because the clustering algorithm used by ADMIXTURE can be influenced by sample order[14] the order of individuals within the 1,804 global cohort was randomized and analyses were run with K in the range from 2-30. Based on the overrepresentation of taurine cattle and domesticated yaks in the 1,804 global cohort, which can bias ancestry assignment[15], taurine cattle and domesticated yak breeds were limited in representation to the 5 individuals with the highest genome sequence coverage, resulting in a cohort of 893 samples with a more balanced breed and species representation. ADMIXTURE analysis was performed on this balanced cohort for K ranging from 2 – 20.

TreeMix
TreeMix-1.13[16] was used to construct a phylogeny rooted at banteng, for populations with at least 3 samples, and that were not known to be breeds formed by the crossing of other breeds, subspecies, or species. The 100K SNP set for these individuals was input to TreeMix to infer migration edges along the phylogeny. Migration edges were added until the increase in the proportion of variation explained by the current model compared to the previous model was less than 0.01.

Subsequent iterations of TreeMix analysis were performed after removing sets of populations that were previously not known to be, but were inferred to be composite, based on ADMIXTURE ancestry assignment and TreeMix edge placement. Redundant patterns of gene flow (*i.e.*, taurine breeds from Western Africa into zebu from Eastern Africa) were simplified through condensing multiple breed level populations from monophyletic and geographically justifiable groupings into a single larger one. This allowed us to preserve these patterns within the context of global *Bos* populations while allowing the software to explore other potential migration edges. Additionally, we iteratively re-ran TreeMix analyses in which the domesticated yaks and gayals with detected domesticated cattle introgression were removed. This left the wild yak, bison, banteng, *Bos frontalis* (gayal from Bangaldesh and India with little to no domesticated introgression), gaur, Chinese wild gaur and aurochs as the only non-domesticated populations. These analyses were performed to test if the detected wild/domesticated admixture originated from strictly wild populations, rather than from potentially detecting domesticated cattle ancestry within populations of domesticated yaks or gayals.

*f*-statistics
To identify populations implicated as possessing more recent admixture and identify the best modern populations representing proxies for the historical donor populations, an $f_3$ test was implemented using AdmixTools-7.0.2[17] qp3Pop. To not introduce bias by preselecting combinations of population, we analyzed every combination of 3 populations (1,898,400 total combinations), where the first two are potential donors and the third is a resulting admixed population, and tests were ranked by their produced Z-scores. The $f_3$ tests with Z-score < -3 were considered significant[17]. Modern populations that were most frequently and significantly predicted to be donors with similar geographic contexts were considered the best ancestral population donor proxies for the predicted admixture events.

Hypotheses of admixture directionality were compared, and ancestry proportions were estimated using the $f_4$-ratio test. For these analyses, models producing a Z-score greater than 3 or less than -3 are considered significant[17], and phylogenies for groups of 5 populations were run, and the test producing the greatest absolute value Z-score was considered to be the best fit.

qpGraph
AdmixTools-7.0.2 qpGraph was used to test hypotheses of admixture detected by TreeMix. Representative breeds for geographically distinct populations were determined based on the best supported donors and recipients from the $f_3$ tests and ancestry clustering from ADMIXTURE. Taurine were represented by a population from Central Asia, Yakut, a population from Central Europe, Braunvieh, and a population from Western Africa, Muturu. Zebu from South Asia were represented by the populations Tharparkar, Indian Zebu, Gir, and Sahiwal. Zebu from East Asia

was represented by Leiqiong Cattle. Wild species used were banteng, the Yenisei aurochs, and a combined population of the wild gaur and Chinese wild gaur.

Demographic models in which a simple tree-like relationship between populations, with a topography based on phylogenies predicted by TreeMix, were fit to represent the null-hypothesis of "no admixture" (Fig. S5a). To first elucidate the role that admixture played in the origin of zebu, gene-flow between gaur and domesticated cattle from the Fertile Crescent (Fig. S5b), or gaur and an aurochs-like ancestor (Fig. S5c) were modeled using qpGraph. A multiple step admixture between aurochs and gaur, then their hybrid (representing Indian aurochs) and domesticated cattle from the Fertile Crescent (Fig. S5d) was also tested. We then incorporated the relationship of taurine from West Africa and zebu from East Asia, either as having wild bovid admixture during their origins, or simple divergence from other populations of domesticated cattle. We created a new null model with all populations having a tree-like relationship (*i.e.*, no admixture). This was comparable to models in which all four of the proposed introgression events were present (Fig. 2) or models where three out of four were included, with the other treated as a simple divergence (Fig. S8).

Multiple network topologies were tested. These include: 1) admixture edge coming from the Yensei 14K branch or the ancestor of auroch and taurine and 2) admixture edge coming from ancestor of gaur/banteng, banteng branch, or gaur branch. Gaur and banteng populations were split into two populations based on NCBI BioProject.

All models were fit using AdmixTools[17] and the R package admixtools2[18], and 10,000 bootstraps were used to test for significant differences in model fits. We also compared the estimated ancestry proportions with ADMIXTURE results, $f_4$-ratio tests, and patterns of ancestry based on published mitochondria and Y haplotype analyses, as well as our own Y chromosome clustering.

Y Chromosome

Sex was estimated for samples using heterozygosity of the X chromosome SNPs and missingness on the Y chromosome and compared to metadata annotations. Samples with missingness >0.9 across the Y chromosome and heterozygosity > 0.01 across the X Chromosome were assigned to be female. Samples with Y missingness < 0.9 and X heterozygosity < 0.01 were assigned to be male. If samples failed both metrics, or agreed but disagreed with annotations from the metadata, sex was assigned "undetermined." Where metrics were unable to determine sex, but metadata were available, the metadata sex was used. All others were assigned "undetermined." Variants from the Y chromosome were extracted for 683 samples determined to be male with genome-wide coverage greater than 10X. These were then filtered for SNP missingness and heterozygous SNPs, resulting in 24,589 SNPs across the Y chromosome. PopART-v1.0[19] was used to generate a network for these markers using the integer NJ net method after 37 samples were excluded due to PopART filters.

cobraa

We used cobraa-v1.1[20] to estimate the timing of predicted introgression events. One sample with the highest genome sequence coverage for each of Sahiwal, Hariana, and Gir was isolated from the 1,804 global cohort VCF, which contained over 135 million variants. For each autosome, input files in the multihetsep format were created, with tandem repeats masked. Tandem Repeats

Finder locations were accessed from the UCSC genome browser (https://hgdownload.soe.ucsc.edu/goldenPath/bosTau9/bigZips/) for the ARS-UCD1.2 genome assembly. All cobraa analyses were performed with the parameter spread_1 set to 0.0003 and spread_2 set to 30. The number of discrete time intervals was left as the default of 32 discrete logarithmic segments (D). The bin size b was also left at the default of 100 and gamma_fg, the initial proportion ancestry from admixture, was set to 0.2. cobraa was set to run until the change in log-likelihood between consecutive iterations was less than 1 using the option –thresh 1 (otherwise, model convergence[20]), or for 400 iterations, whichever came first. A panmictic model (unstructured) was run until convergence was reached for each sample. All possible combinations of divergence time (Te) and reintroduction time (Ts) were performed such that reintroduction time was less than divergence time and both must be less than the 32 time intervals. Structured models that reached convergence were considered to have sufficient estimates for their "goodness of fit" and evaluated against the baseline panmictic model. Comparisons of final log-likelihood scores was used to determine the most likely timing of divergence and reintroduction for each population.

Dsuite Dinvestigate

Using the 38M SNP set, we used Dsuite-v0.5 r57 Dinvestigate[21] to calculate Patterson's D and associated statistics such as $f_{dM}$ for trios of populations using a window of 50 SNPs and a step size of 25 SNPs (average window was approximately 3,250 base pairs). This method uses four populations: 1) an explicitly set donor population, 2) a target population, 3) a population sister to the target, and 4) an outgroup to define ancestral and derived alleles. To investigate introgression from a gaur-like population into *Bos t. indicus* we set Yakut (taurine) as sister to the target population of zebu and used a population combining two wild gaur and eight gayal from Bangladesh and India which clustered with the gaur and show little ancestry from domesticated cattle using ADMIXTURE, with bison as the outgroup (Fig. 3a). Investigation of admixture from taurine into zebu was done setting the population of merged gaurs and gayals as sister to zebu and Yakut as the donor, while using the Yenisei aurochs sample as the outgroup (Fig. 3c). In both cases, the zebu target population was comprised of four cattle breeds from the Indian Subcontinent (Gir, Sahiwal, Hariana, Indian Zebu). We also explored analyses with Braunvieh as the donor of taurine ancestry, and where each of the four zebu breeds from South Asia were treated as individual targets. Windows with $f_{dM}$ in the top 99.5$^{th}$ percentile in the merged zebu and Yakut analyses for both gaur-like and taurine admixture scenarios were collected for functional investigation.

Scanning for $f_{dM}$ along the genome of Muturu, taurine cattle from Western Africa, was done using the Yenisei aurochs as donor of wild ancestry segments and bison as the outgroup, with Braunvieh as sister to Muturu (Fig. S9a). To search for domesticated introgression, Braunvieh, taurine from Central Europe, was used as the donor, with the Yakut taurine breed from North-Central Asia as the outgroup, and Yenisei aurochs as sister to Muturu (Fig. S9c). For the zebu from East Asia, a target population was comprised of two breeds from China, Wannan Cattle and Guangfeng Cattle. Here, searching for wild banteng introgression, an analysis set banteng as the donor, with Gir being the sister to the target, and the population of combined gaurs and gayals as the outgroup (Fig. S10a). A second analysis, looking for domesticated introgression, considered the donor of domesticated ancestry to be Gir, a zebu breed originally from South Asia, with Braunvieh as the outgroup, and banteng as sister to the East Asia cattle (Fig. S10c).

Gene annotation, association, and tissue enrichment analysis
The gProfiler software[22] was used to extract annotated genes (EnsEMBL release 113) within the largest 99.5% of $f_{dM}$ windows. Genes with HGNC gene symbols were input into the Enrichr webtool[23–25]. Enrichment results were retrieved for the ChEA transcription factors 2022, Jensen DISEASES, GO Biological Process 2025, dbGap, UK BioBank GWAS v1, PheWeb 2019, and GWAS Catalog 2023. TissueEnrich[26] was used for tissue enrichment tests. When comparing enriched annotations across South Asia, Africa, and East Asia, a Fisher's combined probability test was calculated for each term.

**Supplementary Text S1**
Candidate genes within and near $f_{dM}$ admixture loci
*Gaur admixture in Bos t. indicus*
Due to the hybrid origin of *Bos t. indicus,* wild introgression has played a major role in differentiating taurine and zebu cattle subspecies and breeds. Fig. 2b shows several peaks of genome-wide $f_{dM}$, which identifies introgression, within which we highlight genes potentially relevant to the specific adaptations of *Bos t. indicus*, particularly in the immune[28] and cardiovascular[29] systems as they relate to pest resistance and heat tolerance. On chromosome 10, *ENSBTAG00000059343* and *ENSBTAG00000059382*, two T-cell receptor genes, are found in a window with large $f_{dM}$ values that likely affect the more resistant immune system of zebu cattle. Chromosome 12 harbors *ABCC4*, which is related to platelet levels[30], and *HS6ST3*, which impacts numerous traits, including body-mass index[31], blood pressure[32] and behavior[33]. Genes *FADS2B*[30] and an *OR5M10*[34] ortholog are also associated with cardiovascular traits and are found within the peak on chromosome 15. Additional immune-related genes are found in peaks on chromosomes 20 and 23. *ANKRD33B* on chromosome 20 is associated with eosinophil levels[35], which may confer the increased pest tolerance of zebu cattle[36,37]. Chromosome 23, the site of the bovine major-histocompatibility complex, had windows with large $f_{dM}$ values which harbored *BOLA-DRB2*, a histocomplex gene, as well as *OR2H18* and *OR2B37*, both of which are tied to BMI[31] and cognitive traits[38] in human GWAS. However, perhaps more importantly in this context, olfactory disfunction is associated with lymphocyte, neutrophil, and basophil cell counts[39]. Thus, the preferential introgression of olfactory receptor genes could provide advantages to the immune system.

*Taurine admixture in Zebu*
Scanning for $f_{dM}$ from the domesticated *Bos t. taurus* donor of ancestry in the origin of *Bos t. indicus* across the genome allows us to identify genes strongly maintained post-hybridization, likely due to their impact on fitness within the domesticated setting. The $f_{dM}$ statistic is sensitive to which populations are set as outgroup and donor, for example switching the placement of Yakut and Yenisei 14K. Thus, we are confident that our analysis has identified genes and genomic regions that reflect the impact of the taurine introgression event, in which domestication related ancestry was conferred to *Bos t. indicus.* Fig. 2d shows more diffuse peaks than in Fig. 2b, but several relevant genes are within these regions. Some of the traits most relevant to domestication are behavior and growth, both of which are highly polygenic. On chromosome 4, we found *THSD7A*, which in human GWAS was tied to cognitive traits[40]. Introgression signal on chromosome 8 pointed to *FREM1*. *FREM1* is associated with BMI[41], cognitive traits[40], and

cardiovascular[42] traits. *SACM1L* on chromosome 22 is associated with cerebral cortex size[43] and cortical thickness[43,44] in human GWAS. These candidate genes point to the important behavioral changes that were necessary for successful domestication.

Other candidate genes highlight other important changes necessary for domestication. On chromosome 4, $f_{dM}$ identified *GIMAP6*, which is associated with total lipids[45], fibrinogen[46] and the blood-clotting response, and lymphocyte count[47].

On chromosome 8, a window with a large $f_{dM}$ statistic, one of the 4 annotated interferon-tau genes, *IFNT3,* was selected in Zebu following introgression from domesticated taurine. In cattle, interferon-tau is responsible for maternal recognition of pregnancy (signal to prevent resumption of estrus cycle), and *IFNT3* is expressed during the critical pregnancy recognition timepoint surrounding day 18 of pregnancy[48]. This locus may have been important in the transition to successful reproduction under human management[49,50] , an essential milestone in domestication[50].

Comparing genes introgessed from Indian aurochs with those introgressed from taurine, only 24 of the 521 genes were shared. Of these 24 genes, only 5 had gene symbols (*ABCA13*, *PRKN*, *TRAV10*, *CFH*, and *CSMD1*). For the remaining 19 genes, two were long noncoding RNAs and 17 were novel protein coding genes from the Ensembl genebuild pipeline, many of which have been retired.

*Aurochs admixture in Africa*

African taurine cattle have elevated $f_{dM}$ signal for aurochs admixture on chromosome 5, which contains a cluster of multiple olfactory receptor genes. While none of these olfactory receptor genes have any associations with traits other than olfaction, the connection between olfactory receptor genes and local adaptation in mammals is abundantly clear[51] and the differentiation in olfactory phenotypes between wild and domestic populations is well documented[52]. Other olfactory receptor genes were found within regions of elevated $f_{dM}$ on chromosome 15, and several of these had an association with cardiovascular traits in human GWAS[53,54]. This includes *OR52E4*[55], *OR52B2*[53], *OR51E1*[54], *ENSBTAG00000007944*[56], *ENSBTAG00000063165*[57], *ENSBTAG00000015876*[58], and *ENSBTAG00000001291*[59].

Multiple genes related to immune system functions were found within regions of aurochs introgression genome-wide. On chromosome 10, *TRAV3,* which was found in human GWAS[60] to impact Chagas disease, caused by trypanosome infection, which is one of the main concerns facing cattle in Africa. *Bos t. taurus* breeds native to West Africa are well known for their trypanotolerance[61]. *TRAV3* affecting trypanosome infection in humans and introgressed into taurine from aurochs provides evidence that trypanotolerance originated in wild African aurochs and was introduced into African taurine via admixture. Figure S9b identifies genes in other immune pathways, such as *ENSBTAG00000059126* and *ENSBTAG00000065000* on chromosome 10, which are T-Cell receptor genes, and *PTPRJ*[30], *P2RX3*[47], *PRG3*[62] and *SLC43A3*[63], which in human GWAS are related to eosinophils[30,47,62,63].

Also separating wild and domesticated animals are pronounced differences in behavioral and cognitive traits[64,65]. Within the loci determined to have been introgressed from aurochs into

African cattle, we found several genes (on chromosome 15: *CTNND1*, *CLP1* and *ZDHHC5*, and on chromosome 24: *SYT4*) that are associated with variation in human behavioral traits[66–69].

Kambal and colleagues[70] reviewed 238 candidate positive selection genes in African cattle populations. Of these only 3 (*FBXO9*, *PCSK6*, and *EPB42*) were identified in our $f_{dM}$ analysis, again supporting the polygenic nature of selection[71] during domestication and post-admixture. *PCSK6* is associated with erythrocyte count and *EPB42* with erythrocyte volume. Anemia is a common symptom of trypanosomiasis infections in African cattle[70], consequently, these genes appear to be strong candidates for trypanosomiasis resilience.

<u>*Taurine admixture in Africa*</u>
Figure S9d shows that genes potentially retained from the domesticated ancestor of African cattle are found throughout the genome, several of which are associated with the hallmark traits of domestication[72,73], including behavior/cognition, reproduction cycle, growth and immunity. *DBI* on chromosome 2, *ZNF330*-like on chromosome 3, *PHLDA1* and *TMTC2* on chromosome 5, *ROCK2* on chromosome 11, and *PGBD5* on chromosome 28 are all associated with behavioral or cognitive traits such as neurodegeneration[74], brain shape [43,75], conduct disorder[76], memory function[77] and brain development[78] in childhood and in aging in human GWAS. The gene *TBC1D22A* was found in humans to be involved in numerous traits, including cortical thickness[79], height[80], hair pigmentation[81], and lifespan[82].

Reproductive genes found within regions of elevated $f_{dM}$ include *GREB1* and *LPIN1*, both of which are found on chromosome 11. Human GWAS found *GREB1* to be associated with female reproductive tract phenotypes[41,83–86] and *LPIN1* with spermatogenesis[30]. Related to reproduction and development within domestic cattle is milk yield, a key component of which is calcium. We found *HPCAL1* also on chromosome 11, which was found to be associated with calcium levels[41]. Finally, genes implicated in the cardiovascular system were detected in this analysis, including *ROCK2*[63,87] and an *RBM17*-like ortholog[87,88]. No genes found in this analysis were also among the candidate genes for selective sweeps in African cattle[70].

<u>*Banteng admixture in East Asia*</u>
Genes found in regions detected as being banteng in origin can be seen in Figure S10b and include *DNAJC10* and *PDE1A* on chromosome 2, and an *HAPLN3* ortholog on chromosome 23. *DNAJC10* is associated with several traits, including growth[80,89], cognition[40], and cardiovascular disease[90]. *PDE1A* is associated with blood pressure[54,58], educational attainment[40], and kidney function[91]. *HAPLN3* on chromosome 23 is associated with growth and overall body size [80,92,93].

<u>*Zebu cattle admixture in East Asia*</u>
Domestic introgression in East Asian zebu cattle shows elevated retention of alleles related to fitness in the domestic niche within the novel environment. Here, we found several genes related to broad spectrum immunity, genome-wide, including the *HSPB9*-like gene on chromosome 1, *GIMAP7* on chromosome 4, and *EXOC3L4* on chromosome 21. *HSPB9* is associated with monocyte levels[63,87], *GIMAP7* is involved in neutrophil, leukocyte and eosinophil counts[30] and triglyceride levels[58], and the *EXOC3L4* gene is linked to platelet volume[92]. Another immune response gene is *CLIC6*, which is involved in inflammatory pathways[94].

Other cardiovascular genes found in this analysis include *GIMAP5*, *GIMAP1* and *IQSEC3*. *GIMAP5* is associated with both blood pressure[59] and pigmentation [95]; another trait often found to be selected in domestication. *GIMAP1* is associated with height[96] and blood pressure[59]. *IQSEC3* is associated with educational attainment[40], body size[80], and blood pressure[54].

**Supplementary Text S2**

Missense mutations in *B. t. indicus* $f_{dM}$ admixture loci

A small number of missense mutations were identified in *B. t. indicus* $f_{dM}$ outlier loci with allele frequencies that matched the history and direction of admixture (Table S7). A missense mutation in *ALPI* originating in the Gaur-like ancestor, which plays a role in immunity in the intestines[97–99], was found in *B. t. indicus* at over a 0.74 frequency, while this mutation is fixed within Gaur and Banteng, over 0.92 in Gayal, but only at 0.08 in *B. t. taurus.* Another missense mutation in *TRMT12* introgressed into *B. t. indicus* from the Gaur-like ancestor is at a frequency of 0.72, but was found to be fixed in nearly all of the analyzed outgroups, and at a frequency of less than 0.03 in *B. t. taurus.* The function of *TRMT12* is poorly understood, but it is associated with height[80] and blood pressure[59] in human.

**Supplementary Text S3**

Aurochs introgression into African *B. t. taurus*

Based on TreeMix migration network results, it has previously been hypothesized that the divergence between African and European *B. t. taurus* cattle was due to admixture between local African aurochs and introduced taurine domesticates[27]. The Yenisei 14K aurochs genome was used to test if African *B. t. taurus* populations have aurochs introgression. Utilizing the $f_4$-ratio test, we estimated aurochs ancestry in West African *B. t. taurus* to be between 19% and 24% (Table S3). Admixture graph modeling indicated that admixture between aurochs and *B. t. taurus* from Central Europe provided a better fit to the population history of *B. t. taurus* in West Africa (Fig. 2, p-value $< 0.0002$) than a model with African *B. t. taurus* being sister to the common ancestor of European *B. t. taurus* (Fig S8a). Aurochs ancestry was estimated to be 11% in West African *B. t. taurus*. No significant $f_3$ statistics were identified for aurochs introgression into West African cattle, indicating that the admixture between aurochs and *B. t. taurus* in Africa was ancient.

**Supplementary Text S4**

Banteng introgression in East Asian cattle

Zebu cattle in Southeast Asia were hybridized with local banteng or Bali cattle (domesticated banteng) populations. ADMIXTURE (Fig. S3) analysis at K=4 found that many of the domesticated cattle breeds from East Asia had between 5 and 20% of their ancestry derived from either gaur or banteng, and this was not detected in any other domesticated population worldwide. TreeMix placed a migration edge from banteng into the root of *B. t. indicus* in East Asia after its split from the South Asian *B. t. indicus* lineage (Fig. S4). The $f_3$ analyses found the Jian and Leiqiong cattle breeds from Southern China to have significant signals of introgression (Table S2) from banteng and Dehong (a *B. t. indicus* population from western China, which was most similar to *B. t. indicus* from South Asia in the ADMIXTURE analyses). Additionally, the $f_3$ statistics indicated that Leiqiong was the most frequent and significant donor population of *B. t.*

*indicus* ancestry for the hybrid domesticated cattle populations in East Asia (Table S2), reinforcing the importance of banteng ancestry in the cattle of this region. Several $f_4$-ratio tests found banteng ancestry in East Asian *B. t. indicus* to be between 14% and 35% (Table S3), with the most strongly supported estimate being 18.7% banteng ancestry. In the admixture graph analyses, a model which allowed introgression from banteng, which was estimated at 26% ancestry (Fig 1e) performed substantially better (p-value < 0.0002, Table S4) than a model with East Asian *B. t. indicus* (Leiqiong cattle) as a non-admixed sister group to *B. t. indicus* from South Asia (Fig. S8b). Unlike the *Bos gaurus* introgression into the ancestors of South Asian *B. t. indicus*, for which no ADMIXTURE or $f_3$ signal for recent introgression was detected, the *Bos javanicus* introgression into East Asian *B. t. indicus* was identified by the ADMIXTURE and $f_3$ statistics. Thus, the *Bos javanicus* introgression into East Asian *B. t. indicus* breeds is a much more recent event than the admixture event that formed *B. p. namadicus*.

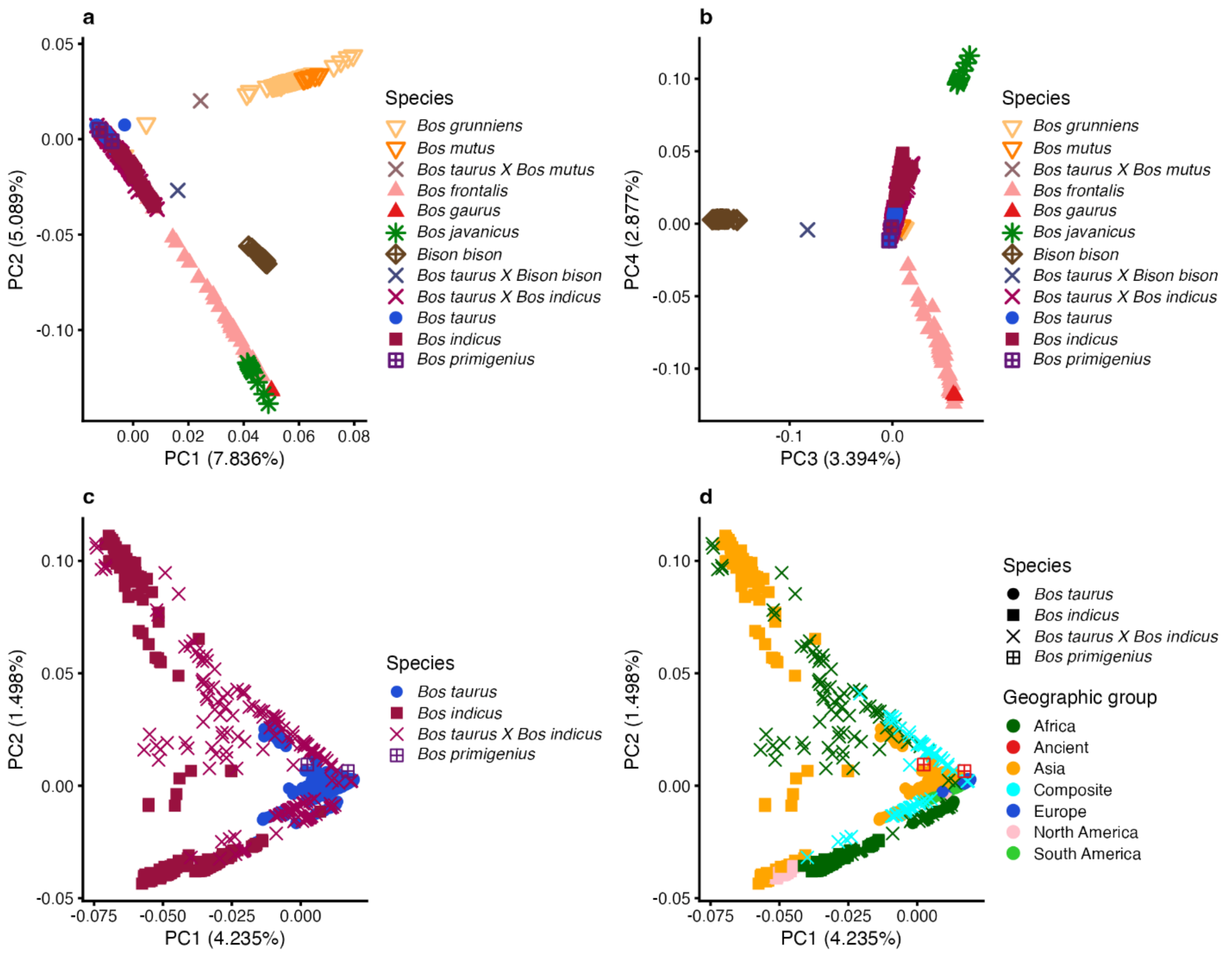


**Fig. S1.**

**Principal Component Analysis.** (**A**) Principal components 1 and 2 for 1,804 Bovids (**B**) Principal components 3 and 4 for 1,804 Bovids (**C**) Principal components 1 and 2 for 1,531 domesticated cattle and aurochs (**D**) Principal components 1 and 2 for 1,531 domesticated cattle and aurochs. Analysis based on 38,097,187 linkage disequilibrium pruned SNPs. Color indicates historical geographic region, with the exception of "Ancient", denoting the aurochs samples, and "Composite" denoting recently developed composite breeds with ancestry from multiple geographic regions.

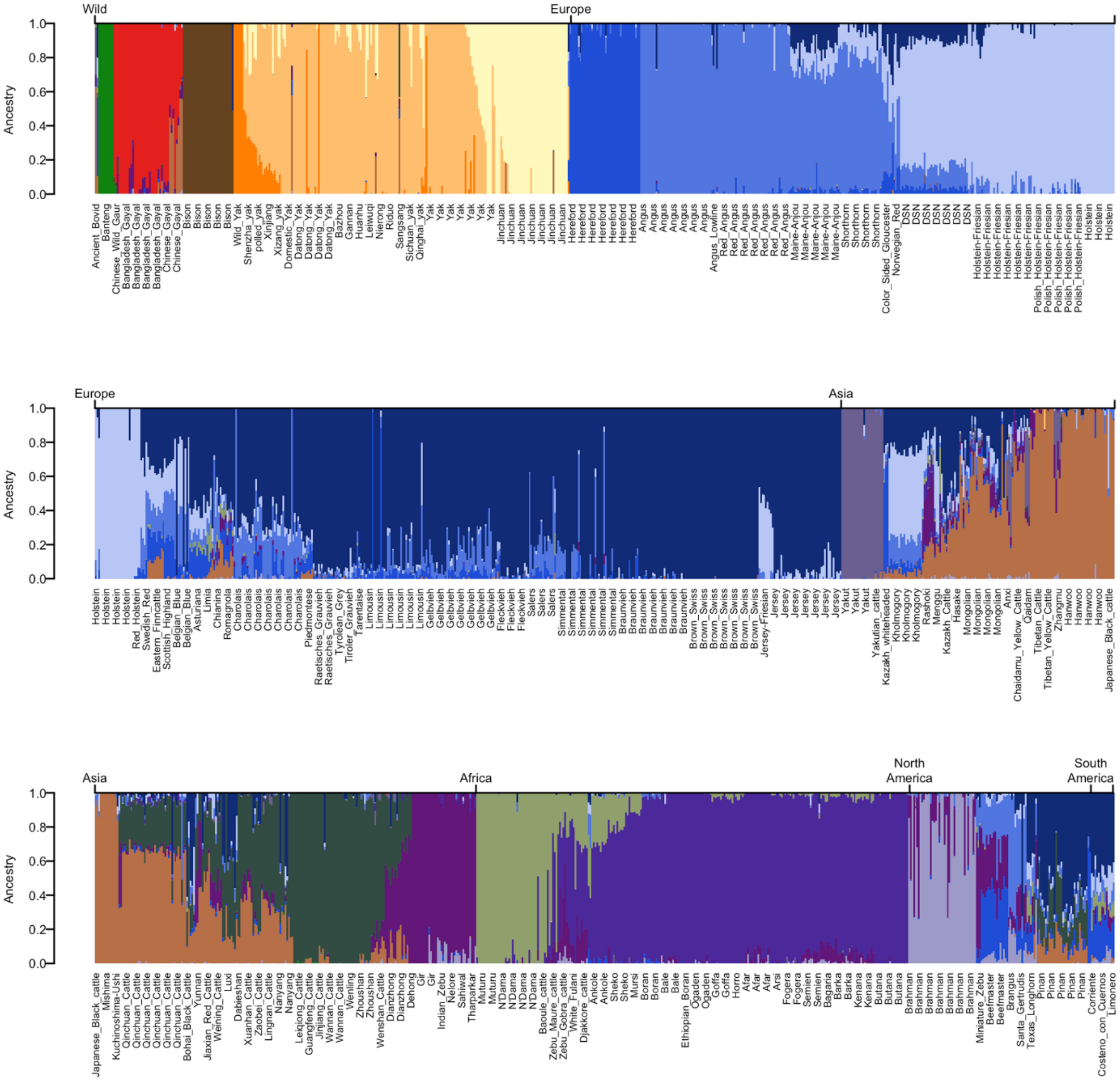


**Fig. S2.**

**Ancestry clustering using ADMIXTURE for 1,804 bovids with K=17 based on 100,000 randomly sampled SNPs.** Samples are arranged in order of species, broad historical geographic placement, then breed. Note that the three panels represent one continuous plot read left to right.

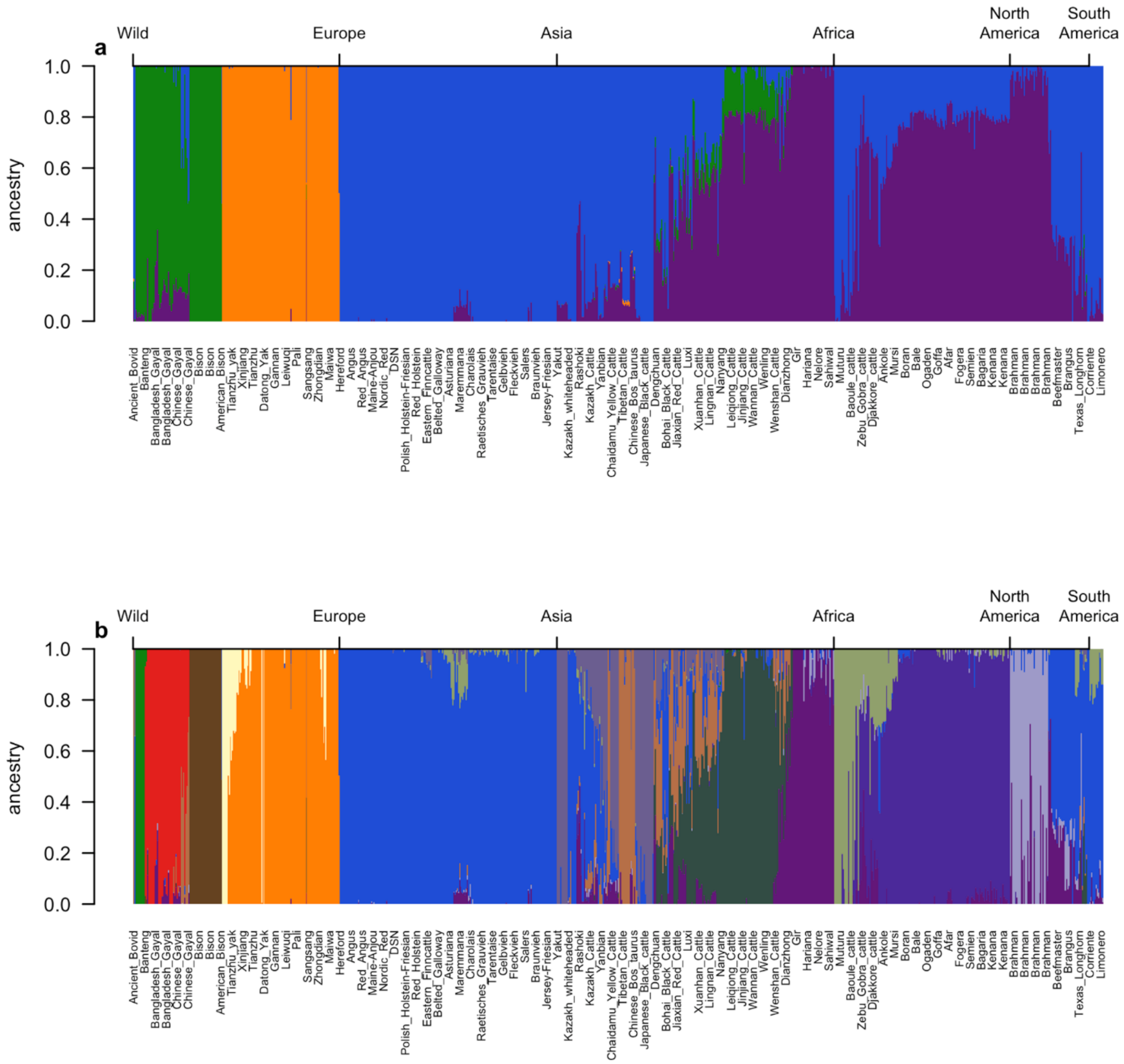


**Fig. S3.**

**Ancestry clustering using ADMIXTURE for 893 bovids. (A)** Model with K = 4, using 100,000 randomly sampled SNPs. **(B)** Model with K=13 based on 100,000 SNPs. Samples are arranged in order of species, broad historical geographic placement, then breed.

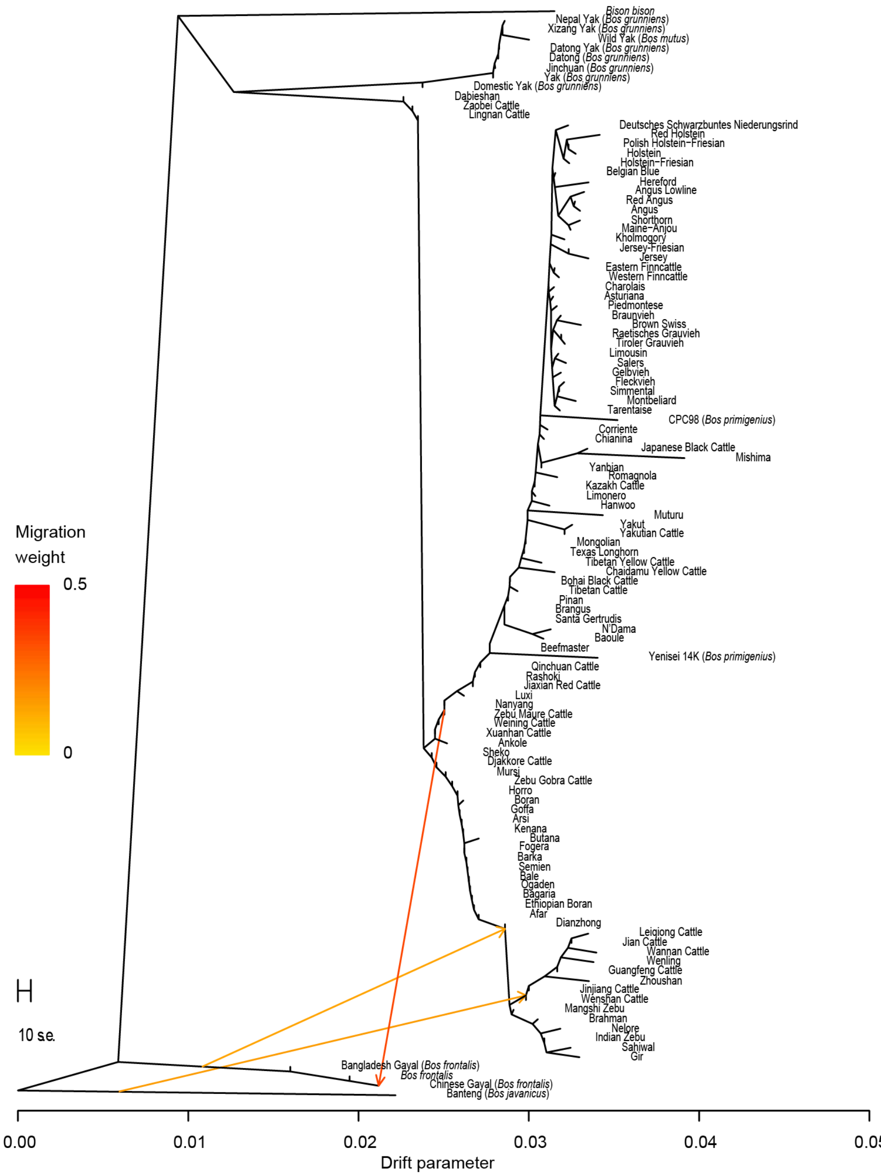


**Fig. S4.**

**TreeMix network of Bovid populations**. Analysis included only populations with at least 3 samples, except for the two ancient samples, and was based on 100,000 randomly sampled SNPs. Populations known *a priori* to be admixed (*e.g.*, from known breed formation information) were excluded to allow the network to explore unknown migration edges. The first edge added to the network represented gene flow between banteng and the root of zebu from East Asia (ancestry proportion = 0.157, p-value < 2.2e-308), and the migration edge between the gaur and gayal branch into the root of zebu was the second added edge (ancestry proportion = 0.144, p-value < 2.2e-308).

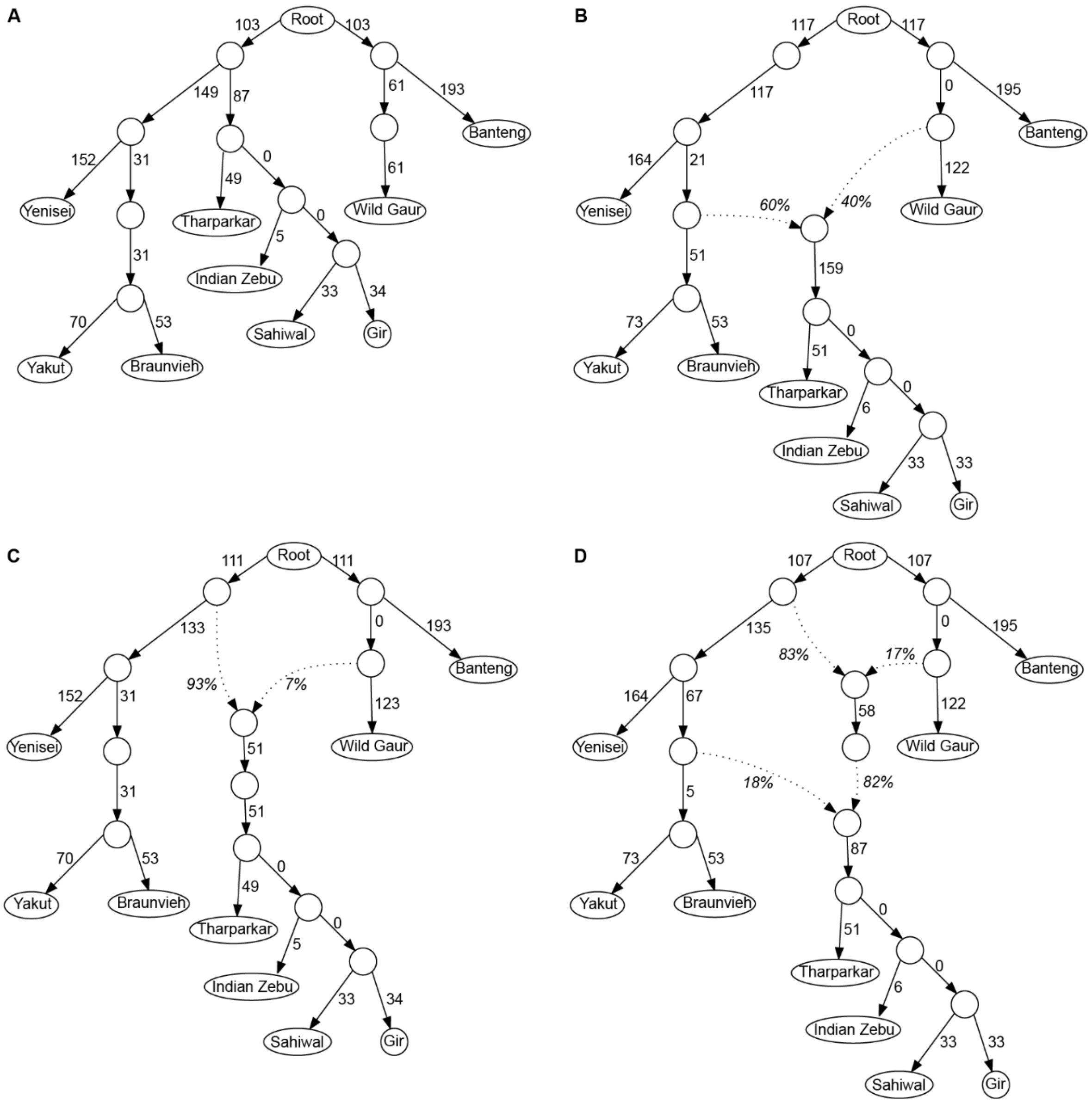


**Fig. S5.**

**Admixture graphs created with qpGraph for South Asian zebu (*Bos taurus indicus*).** Graph modeling **(A)** A bifurcating phylogenetic relationship between taurine, *Bos primigenius* from Yenisei, zebu, *Bos javanicus* and *Bos gaurus* with no admixture. The phylogeny is based on TreeMix phylogenetic networks without admixture. **(B)** Gene flow between the domesticated cattle from the Fertile Crescent and a gaur-like ancestor resulting in the origin of zebu. **(C)** Gene flow from a gaur-like ancestor into the progenitor of zebu, Indian aurochs (*Bos p. namadicus*). This network accounts for the gaur introgression detected in TreeMix, without gene flow from the previously domesticated taurine from the Fertile Crescent. Zebu is then independently domesticated from Indian aurochs. **(D)** Older admixture from a gaur-like ancestor to produce Indian aurochs, followed by a subsequent admixture with domesticated taurine from the Fertile Crescent to produce zebu.

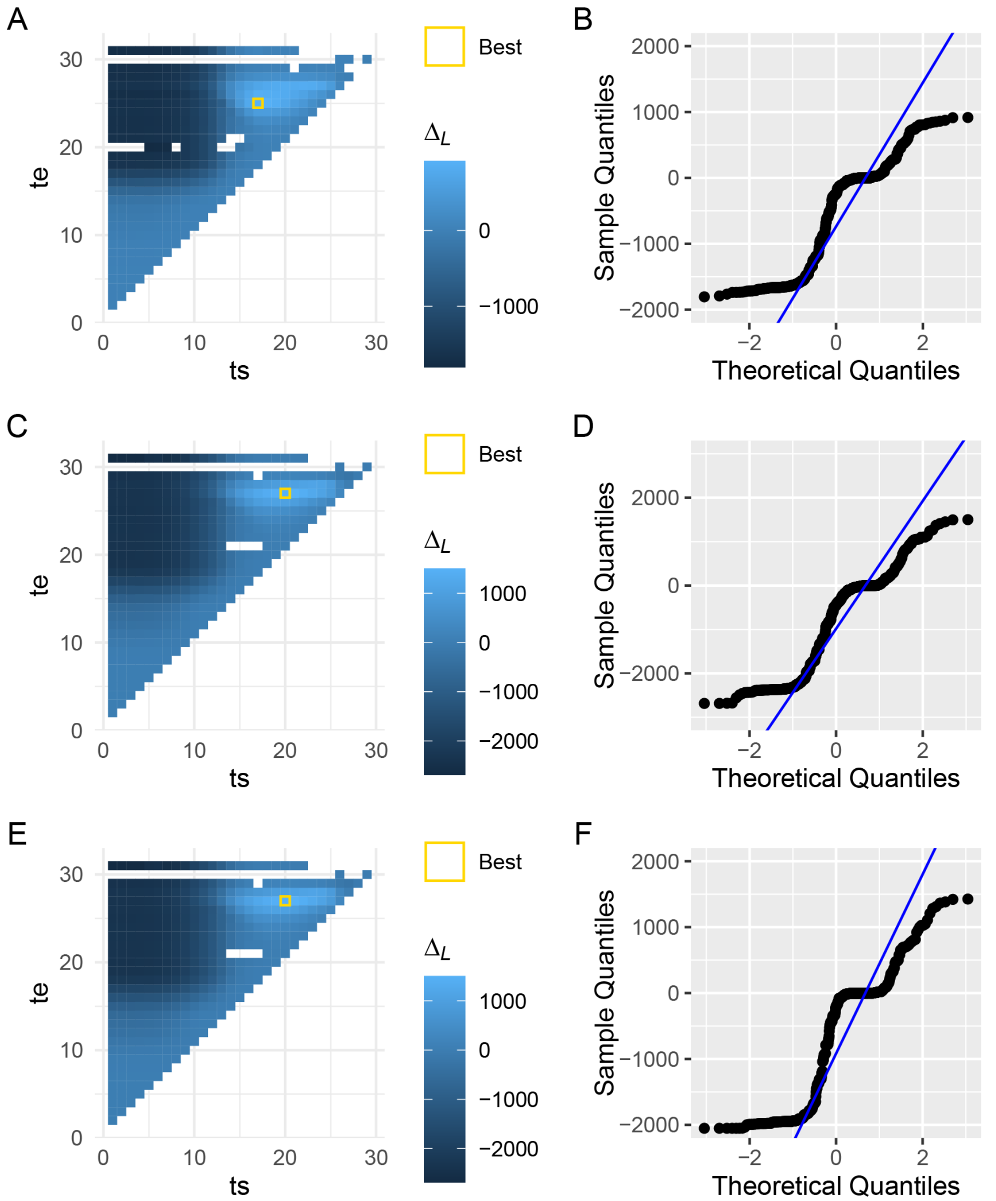


**Fig. S6. Difference in log likelihood ($\Delta_L$) between panmictic and structured models and quantile-quantile plot of those differences.** A) Heatmap of $\Delta_L$ for highest coverage Sahiwal genome, where "te" is the discrete time interval window of the population split time (T2) and "ts" is the population admixture time (T1). B) Quantile-quantile plot of the $\Delta_L$ scores of the various models for Sahiwal. C) Heatmap of $\Delta_L$ for highest coverage Gir genome. D) Quantile-quantile plot of the $\Delta_L$ scores of the various models for Gir. E) Heatmap of $\Delta_L$ for highest coverage Hariana genome. F) Quantile-quantile plot of the $\Delta_L$ scores of the various models for Hariana.

.

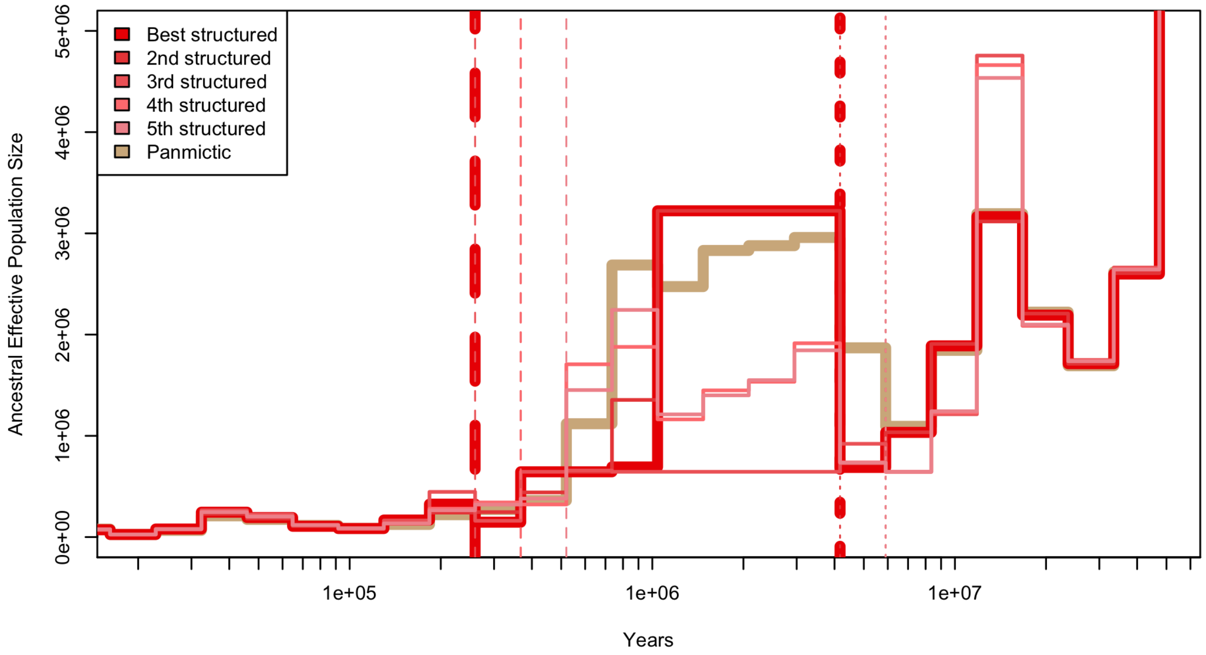


**Fig. S7.**

**Coalescence based estimates of changes in effective population size and the timing of admixture from an unsampled ghost population for the *Bos taurus indicus* breed Sahiwal**. The tan line represents this populations estimated history under a "panmictic" model, otherwise known as an unstructured, or no admixture model. The red toned lines represent the same population, but under a model that presumes a historic divergence (split) of the population into the focal population and an unsampled ghost population, with a subsequent admixture event (rejoin) between the two, otherwise known as structured models. The vertical dashed lines on the right side of the graph indicate the split tine of the ghost population, and the vertical dotted lines on the left side of the plot represent the rejoin time. Gamma (admixture proportion) is 0.465 (range of top models from 0.298 to 0.702). A generation interval of 5 years and a mutation rate of 1.2e-8 was used to convert from coalescent units to years.

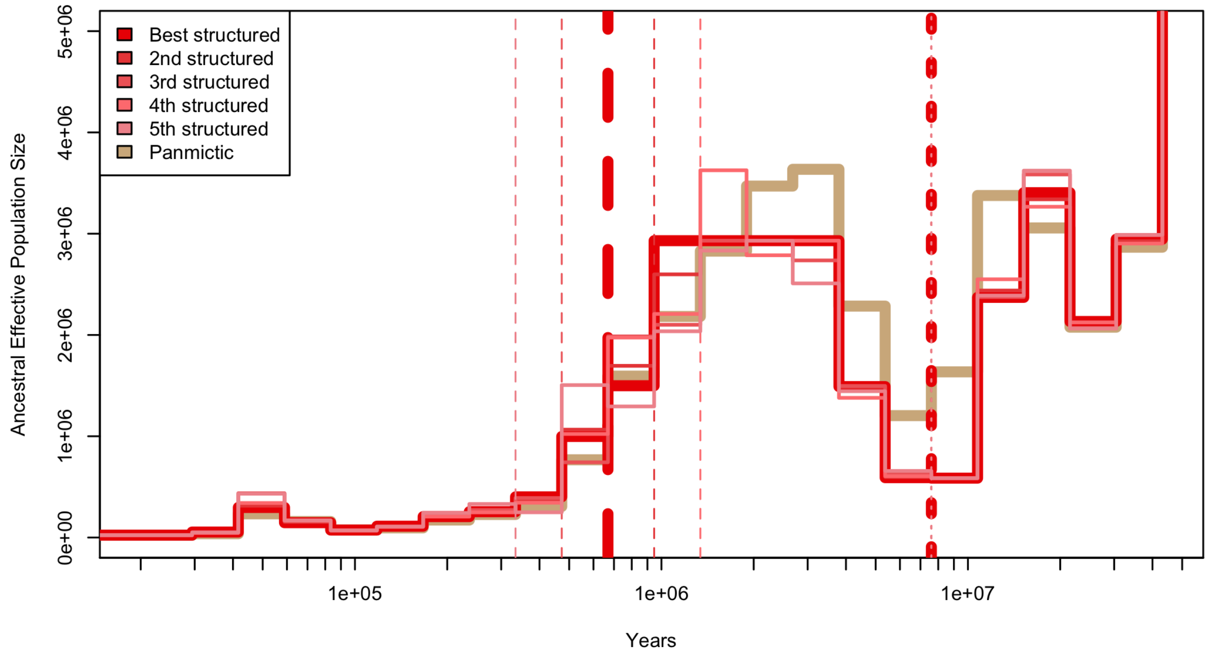


**Fig. S8.**

**Coalescence based estimates of changes in effective population size and the timing of admixture from an unsampled ghost population for the *Bos taurus indicus* breed Gir**. The tan line represents this populations estimated history under a "panmictic model," otherwise known as an unstructured, or no admixture model. The red toned lines represent the same population, but under a model that presumes a historic divergence (split) of the population into the focal population and an unsampled ghost population, with a subsequent admixture event (rejoin) between the two, otherwise known as structured models. The vertical dashed lines on the right side of the graph indicate the split tine of the ghost population, and the vertical dotted lines on the left side of the plot represent the rejoin time. Gamma (admixture proportion) is 0.280 (range of top models from 0.252 to 0.280). A generation interval of 5 years and a mutation rate of 1.2e-8 was used to convert from coalescent units to years.

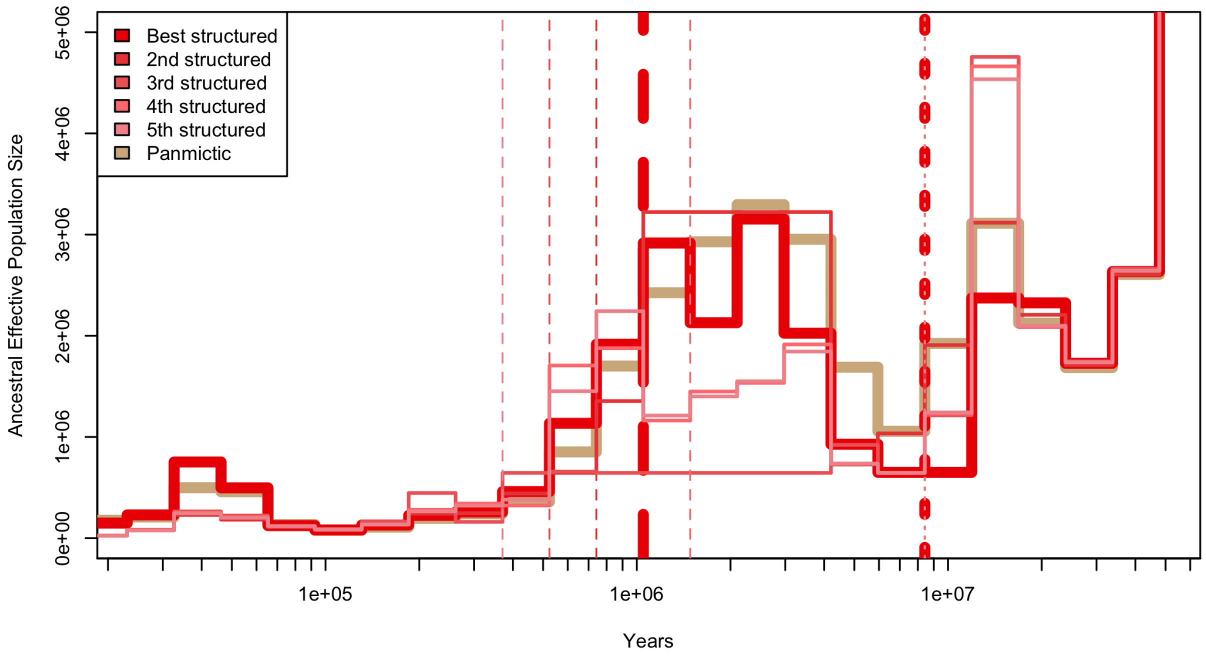


**Fig. S9.**

**Coalescence based estimates of changes in effective population size and the timing of admixture from an unsampled ghost population for the *Bos taurus indicus* breed Hariana.** The tan line represents this populations estimated history under a "panmictic model," otherwise known as an unstructured, or no admixture model. The red toned lines represent the same population, but under a model that presumes a historic divergence (split) of the population into the focal population and an unsampled ghost population, with a subsequent admixture event (rejoin) between the two, otherwise known as structured models. The vertical dashed lines on the right side of the graph indicate the split tine of the ghost population, and the vertical dotted lines on the left side of the plot represent the rejoin time. Gamma (admixture proportion) is 0.223 (range of top models from 0.213 to 0.224) A generation interval of 5 years and a mutation rate of 1.2e-8 was used to convert from coalescent units to years.

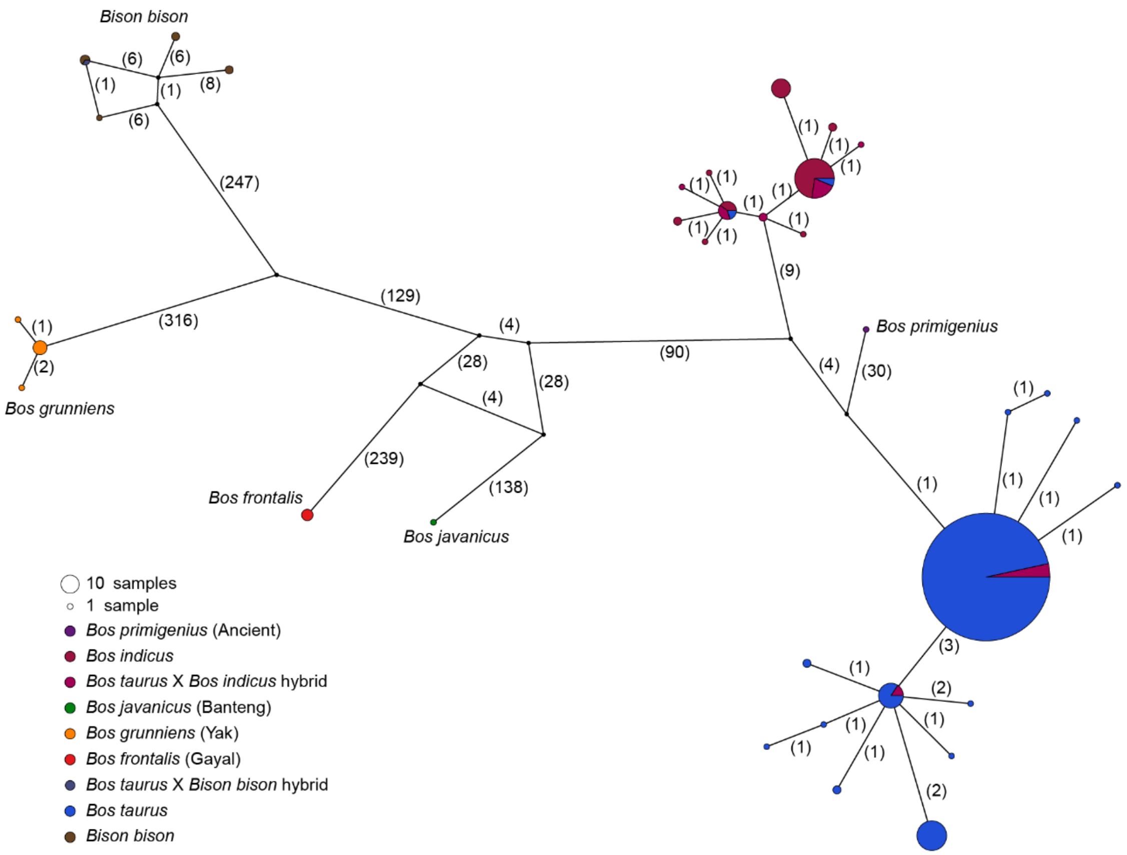


**Fig. S10.**

**Network of Y chromosome haplotypes.** Network for 683 male samples with genome-wide coverage > 10X. Filtering for non-heterozygous sites and no missingness resulted in 24,589 Y chromosome SNPs. Numbers in parentheses represent the number of nucleotide differences between nodes.

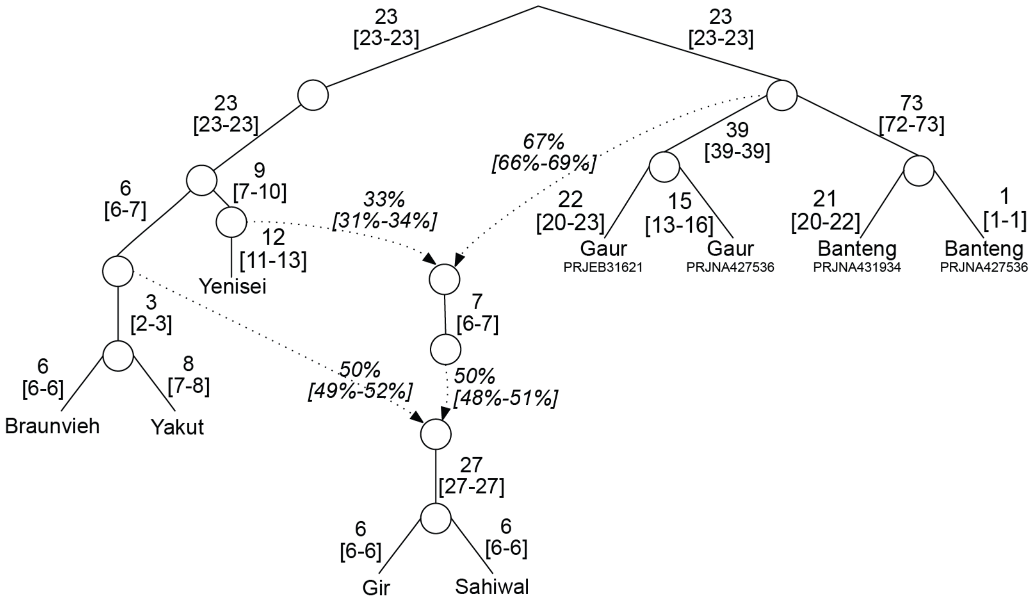


**Fig. S11.**

**Admixture graph modeling older admixture from a wild *Bos* ancestor and the branch leading to the Yenisei 14K sample to produce Indian aurochs, followed by a subsequent admixture with domesticated taurine from the Fertile Crescent to produce zebu.** Because aurochs ancestry is less than 50%, this graph is not congruent with Y chromosome and mtDNA data. Score for this graph was 246.678.

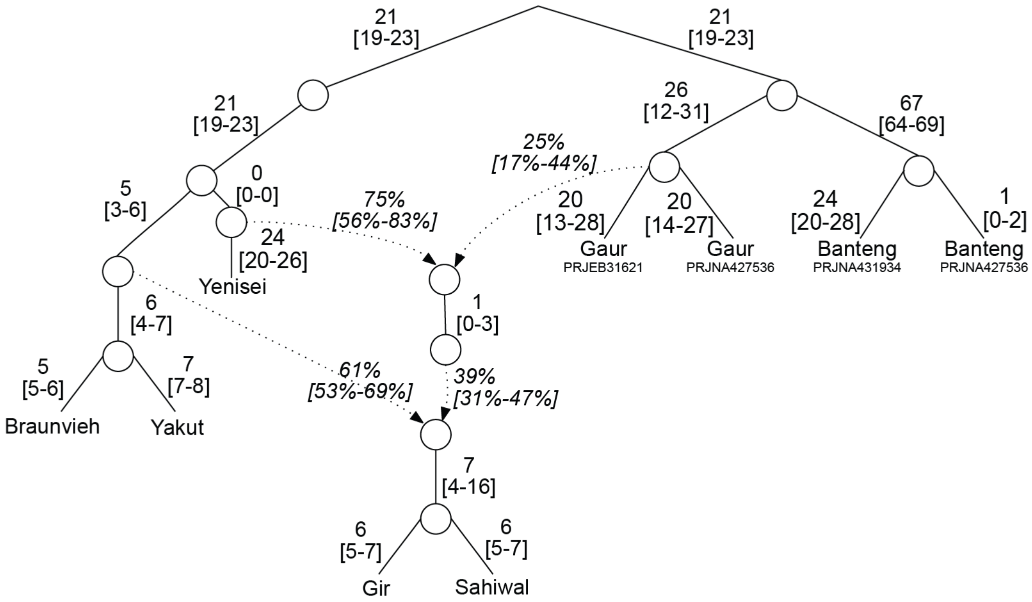


**Fig. S12.**
**Admixture graph modeling older admixture from a gaur ancestor and the branch leading to the Yenisei sample to produce Indian aurochs, followed by a subsequent admixture with domesticated taurine from the Fertile Crescent to produce zebu.** Score for this graph was 21061.700.

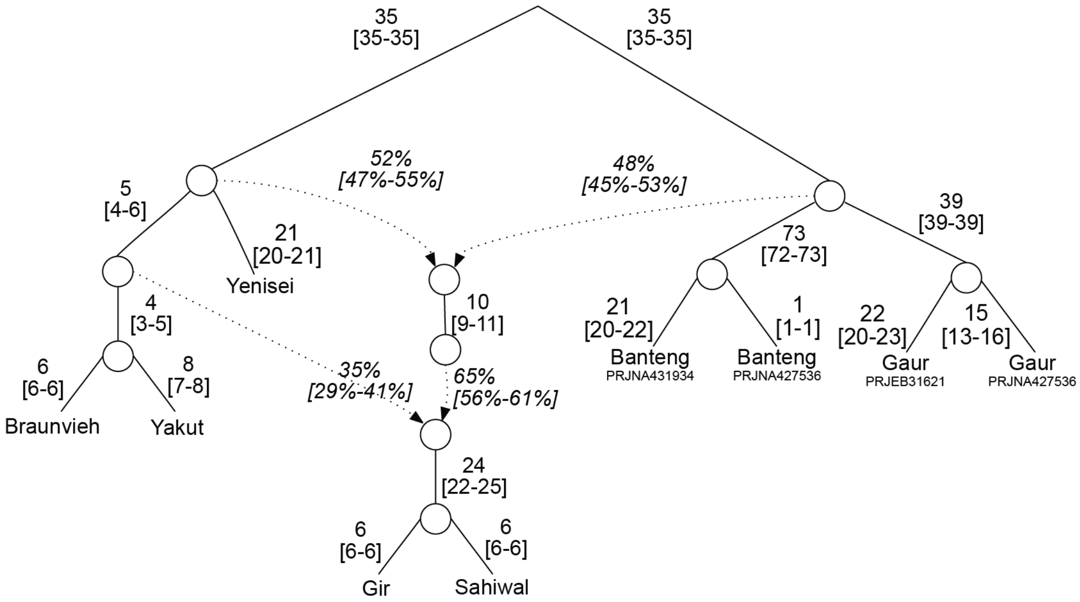


**Fig. S13.**
**Admixture graph modeling first admixture from node ancestral to gaur and banteng and the node ancestral to aurochs and taurine cattle to produce Indian aurochs, followed by a subsequent admixture with domesticated taurine from the Fertile Crescent to produce zebu.** Because aurochs ancestry is nearly 50%, this graph is not likely given Y chromosome and mtDNA data. Score for this graph was 246.911.

.

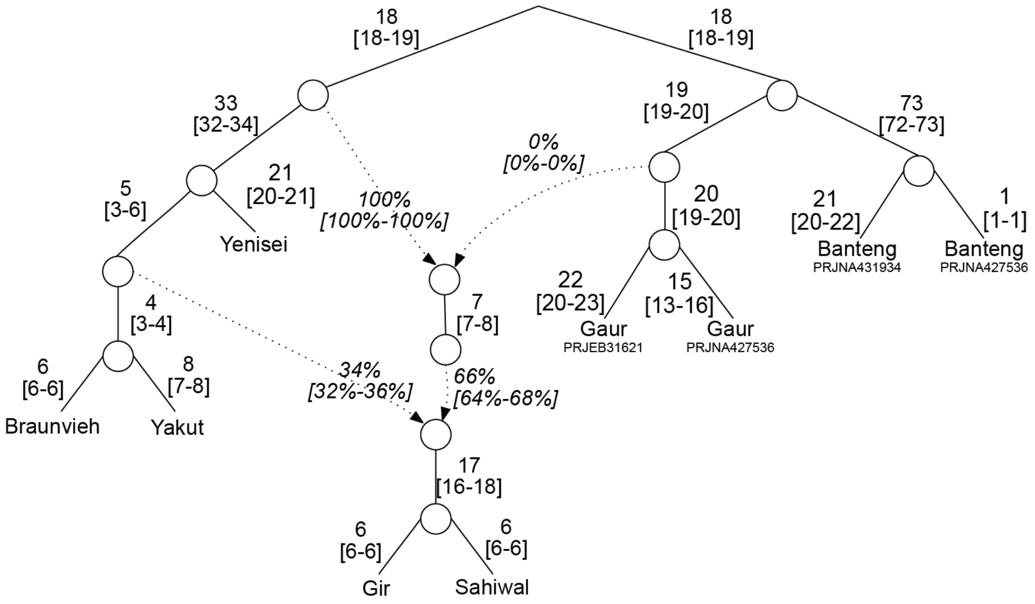


**Fig. S14.**
**Admixture graph modeling older admixture from an older gaur ancestor and an edge ancestral to aurochs and taurine cattle to produce Indian aurochs, followed by a subsequent admixture with domesticated taurine from the Fertile Crescent to produce zebu.** Note zero ancestry from wild *Bos* edge. Score for this graph was 246.896.
.

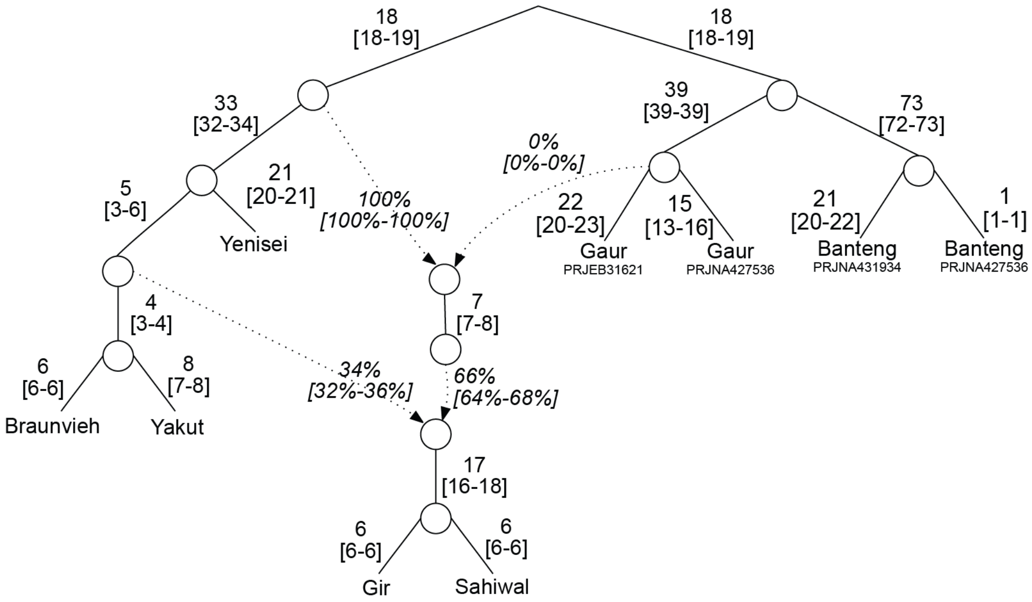


**Fig. S15.**
**Admixture graph modeling first admixture from a gaur ancestor and an edge ancestral to aurochs and taurine cattle to produce Indian aurochs, followed by a subsequent admixture with domesticated taurine from the Fertile Crescent to produce zebu.** Note zero ancestry from wild *Bos* edge. Score for this graph was 247.340.
.

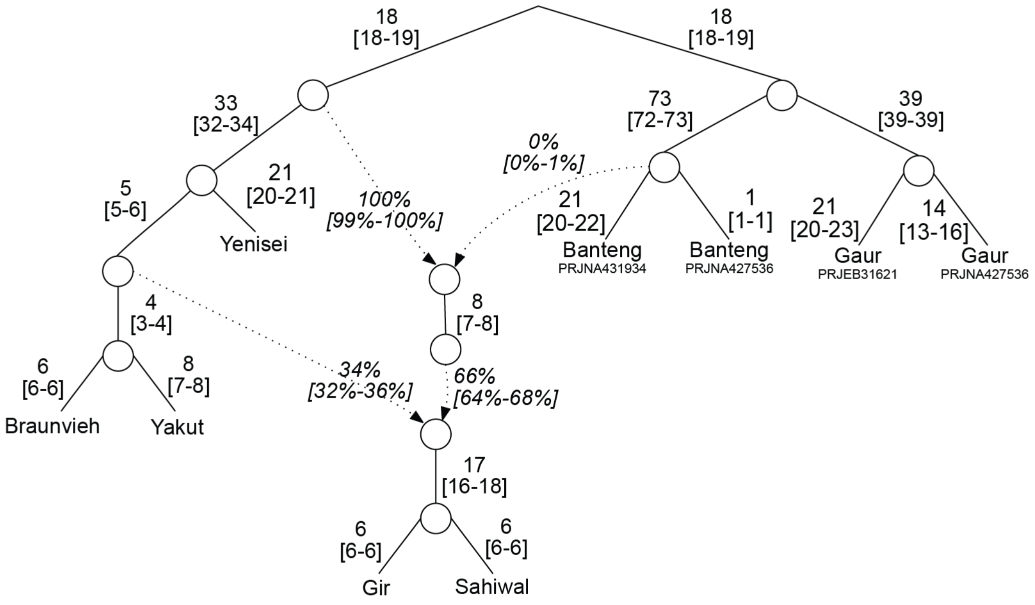


**Fig. S16.**
**Admixture graph modeling first admixture from a banteng ancestor and an edge ancestral to aurochs and taurine cattle to produce Indian aurochs, followed by a subsequent admixture with domesticated taurine from the Fertile Crescent to produce zebu.** Note zero ancestry from wild *Bos* edge. Score for this graph was 202.663.

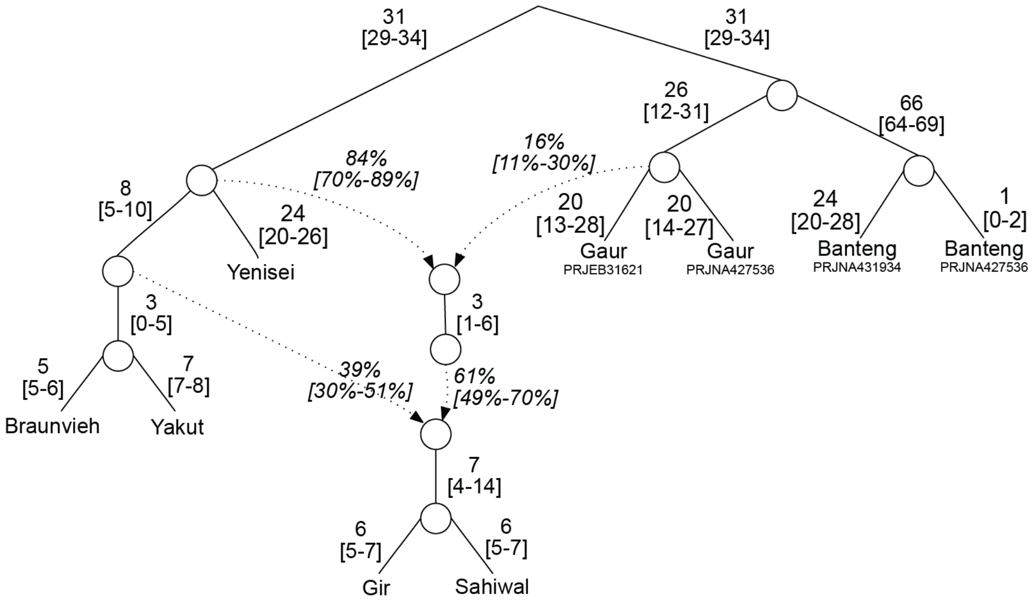


**Fig. S17.**
**Admixture graph modeling first admixture from gaur ancestral node and ancestral aurochs-taurine node to produce Indian aurochs, followed by a subsequent admixture with domesticated taurine from the Fertile Crescent to produce zebu.** Score for this graph was 21064.597.

.

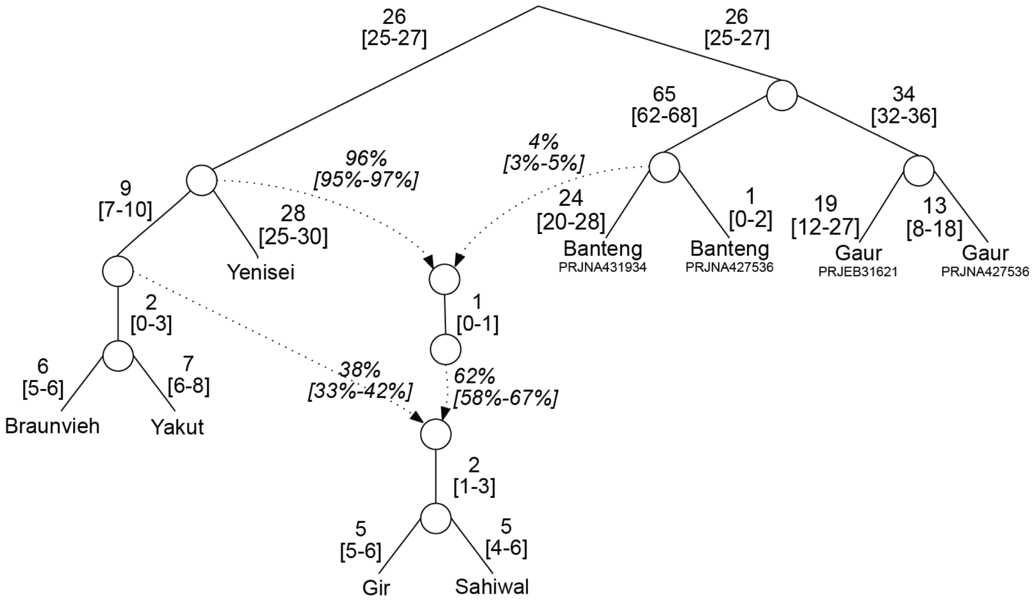


**Fig. S18.**
**Admixture graph modeling first admixture from banteng ancestral node and ancestral aurochs-taurine node to produce Indian aurochs, followed by a subsequent admixture with domesticated taurine from the Fertile Crescent to produce zebu.** Score for this graph was 23675.686.

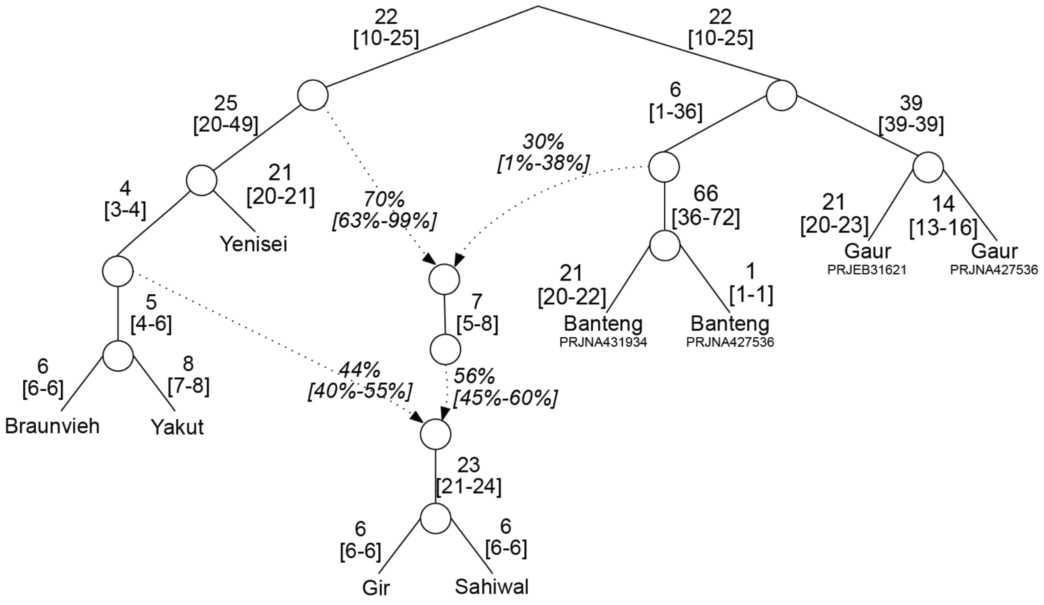


**Fig. S19.**
**Admixture graph modeling first admixture from banteng ancestral edge and ancestral aurochs edge to produce Indian aurochs, followed by a subsequent admixture with domesticated taurine from the Fertile Crescent to produce zebu.** Note short branch length from wild *Bos* ancestor to admixture node paired with long branch length from admixture to banteng node. Score for this graph was 202.372.

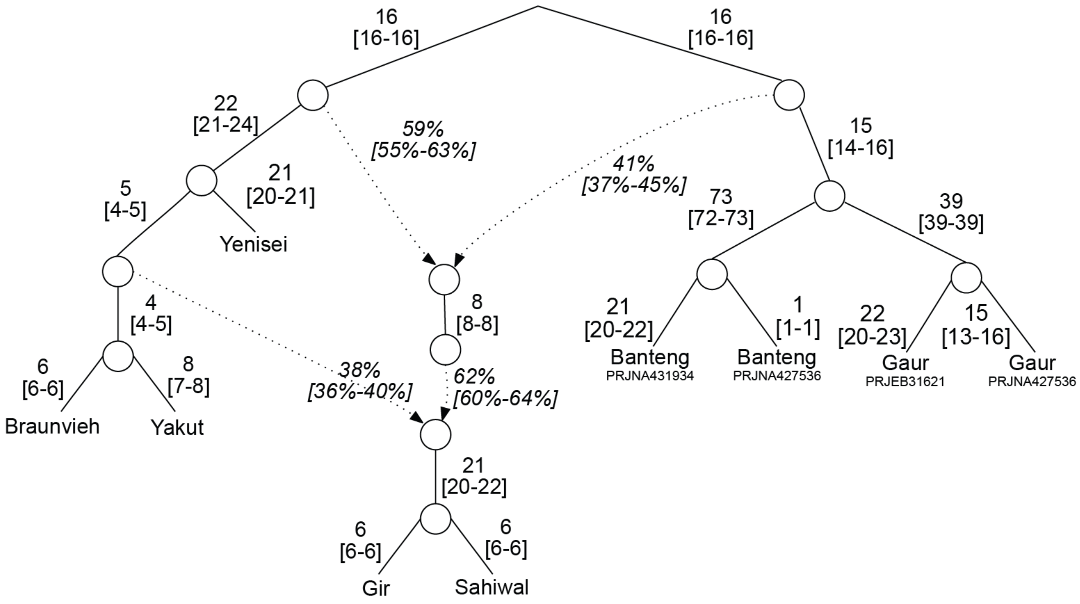


**Fig. S20.**
**Admixture graph modeling first admixture from wild *Bos* ancestral edge and ancestral aurochs edge to produce Indian aurochs, followed by a subsequent admixture with domesticated taurine from the Fertile Crescent to produce zebu.** Score for this graph was 246.898.
.

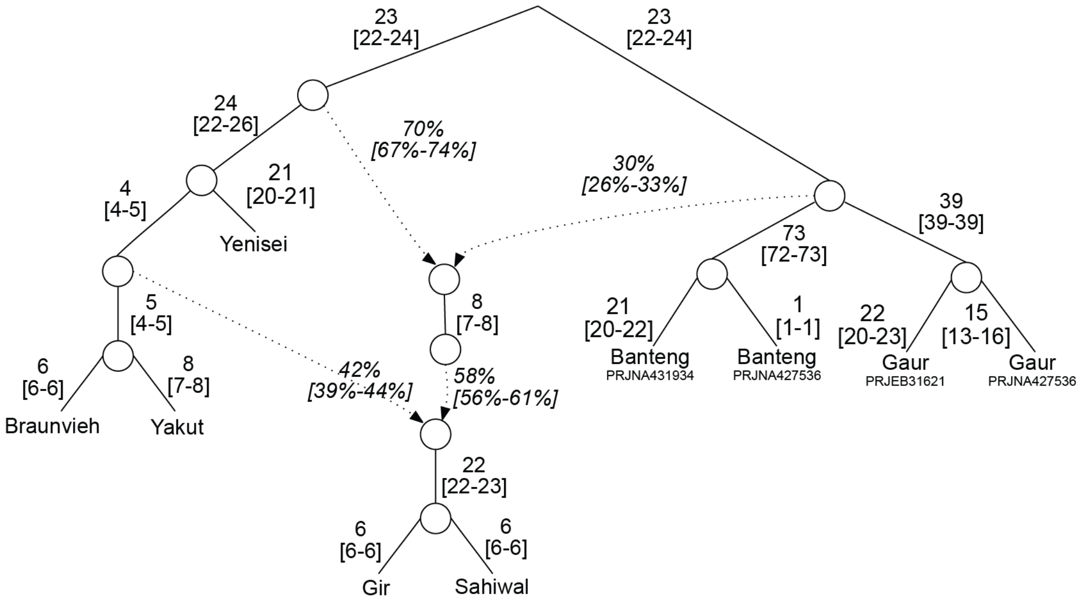


**Fig. S21.**
**Admixture graph modeling first admixture from wild *Bos* ancestral node and ancestral aurochs edge to produce Indian aurochs, followed by a subsequent admixture with domesticated taurine from the Fertile Crescent to produce zebu.** Score for this graph was 247.335.

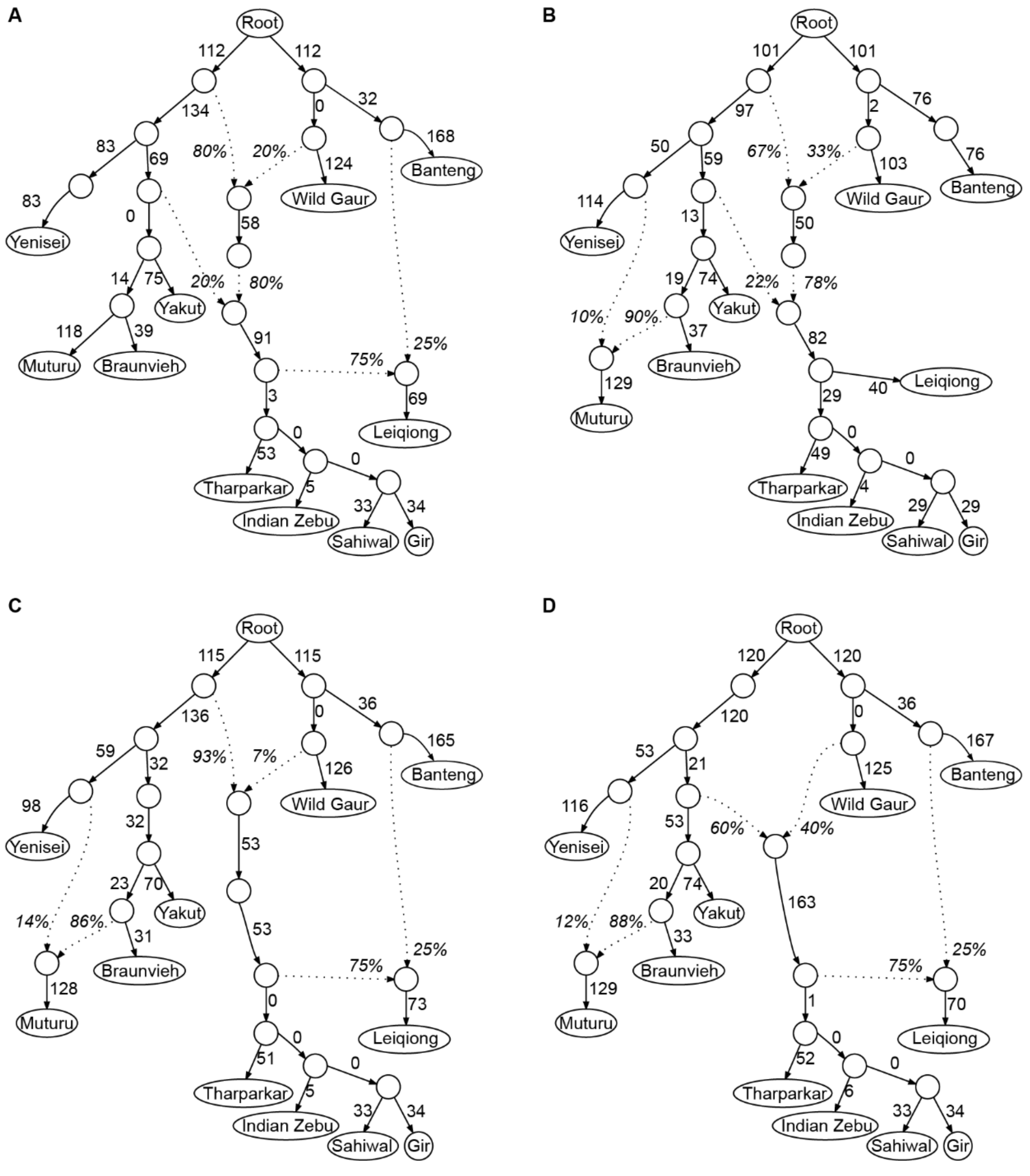


**Fig. S22.**

**Admixture graphs created with qpGraph for South Asia, Africa, and East Asia admixture events. (A)** A multi-step admixed origin of zebu as well as an admixed origin of East Asian zebu cattle, but a non-admixed origin of African taurine cattle. **(B)** A multi-step admixed origin of zebu as well as an admixed origin of African taurine cattle, but a non-admixed origin of East Asian zebu cattle. **(C)** Admixture from a gaur-like ancestor into Indian aurochs with a subsequent independent domestication of zebu as well as an admixed origin of East Asian zebu cattle, and an admixed origin of African taurine cattle. **(D)** Admixture between a gaur-like ancestor and domesticated taurine cattle from the Fertile Crescent producing zebu as well as an admixed origin of East Asian zebu cattle, and an admixed origin of African taurine cattle.

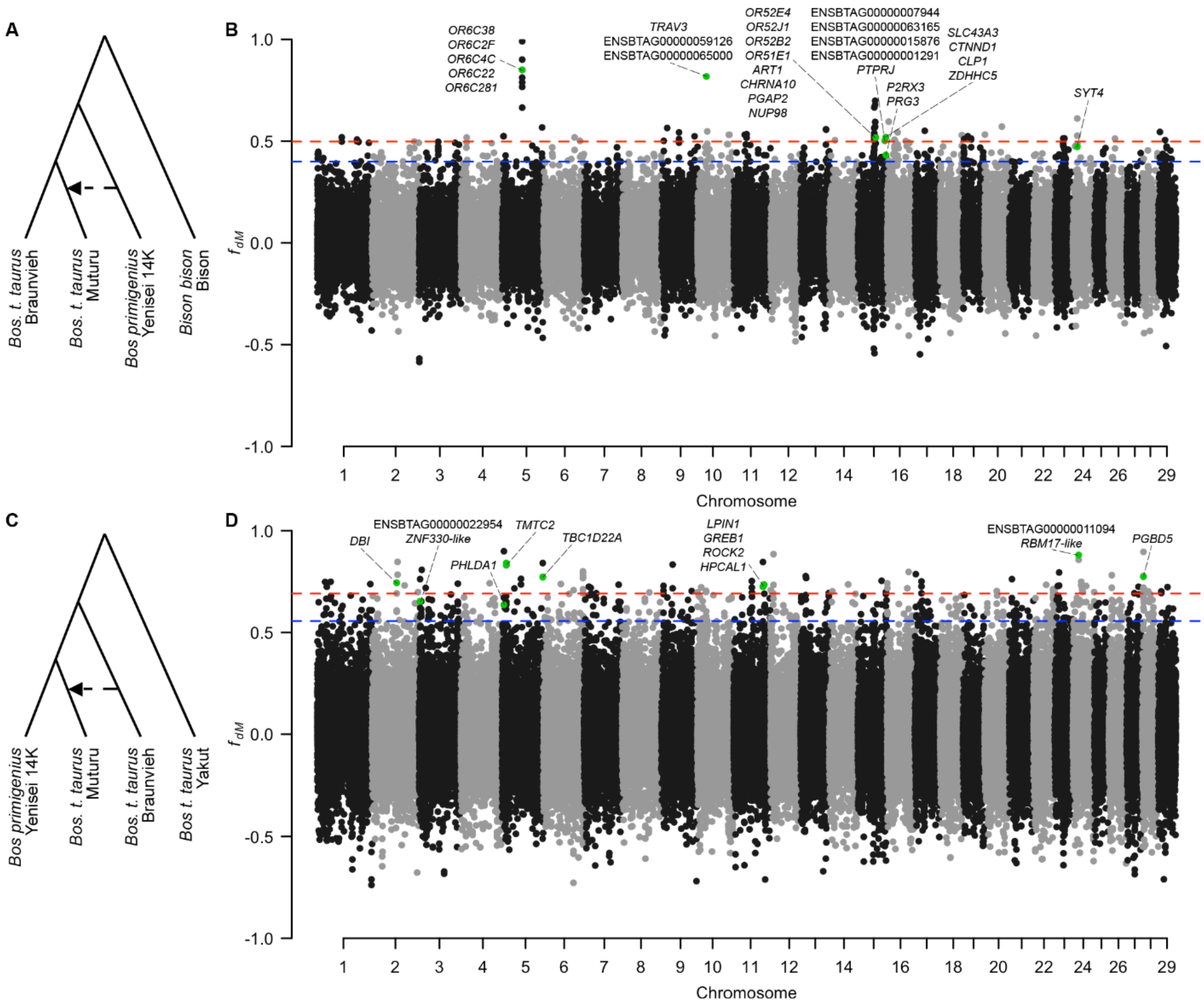


**Fig. S23.**

**Genome-wide scans of $f_{dM}$ statistics for African *Bos taurus taurus* cattle (Muturu) using Dsuite Dinvestigate. (A)** Phylogeny and populations used in $f_{dM}$ analysis of wild introgression into taurine in Africa. **(B)** Manhattan plot of $f_{dM}$ values for wild introgression into taurine in Africa. **(C)** Phylogeny and populations used in the $f_{dM}$ analysis of domesticated cattle introgression into taurine in Africa. **(D)** Manhattan plot of $f_{dM}$ values for domesticated introgression into taurine in Africa. Dashed blue line represents the 99.5$^{th}$ percentile threshold and the dashed red line indicates the 99.9$^{th}$ percentile threshold. Loci above the 99.5$^{th}$ percentile were used to investigate enrichment. Loci containing discussed candidate genes are plotted in green.

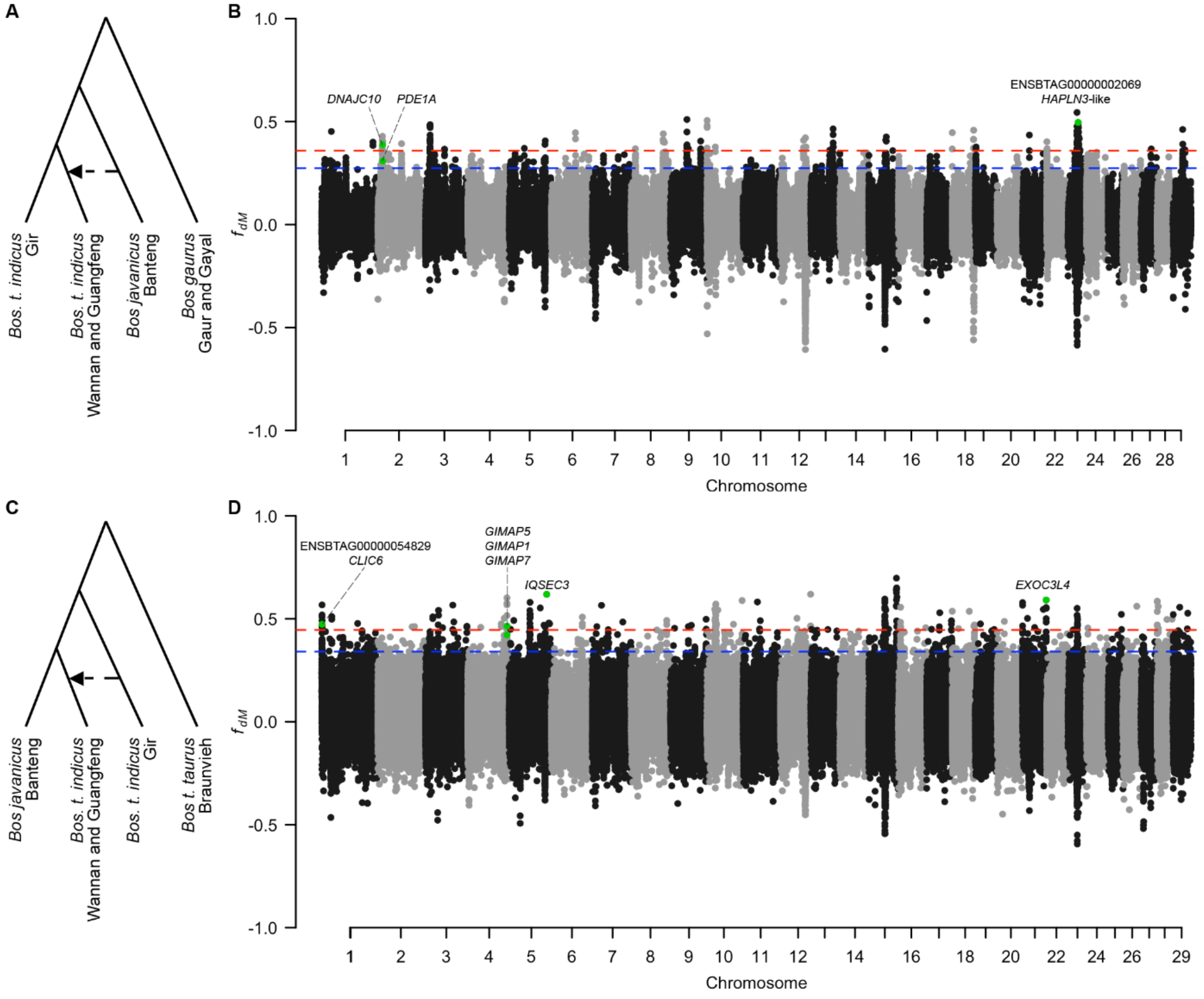


**Fig. S24.**

**Genome-wide scans of *$f_{dM}$* statistics for *Bos taurus indicus* from East Asia, using Dsuite Dinvestigate. (A)** Phylogeny and populations used in $f_{dM}$ analysis of Banteng introgression into zebu in East Asia. **(B)** Manhattan plot of $f_{dM}$ values for Banteng introgression into zebu in East Asia. **(C)** Phylogeny and populations used in the $f_{dM}$ analysis of domesticated cattle introgression into zebu in East Asia. **(D)** Manhattan plot of $f_{dM}$ values for domesticated introgression into zebu in East Asia. Dashed blue line represents the 99.5$^{th}$ percentile threshold and the dashed red line indicates the 99.9$^{th}$ percentile threshold. Loci above the 99.5$^{th}$ percentile were used to investigate enrichment. Loci containing discussed candidate genes are plotted in green.

**Table S1. (separate file)**
Sample provenance, species, geographic, breed and sex information, and sequencing coverage.

**Table S2. (separate file)**
Largest $f_3$ statistic for those populations with significant $f_3$ scores.

**Table S3. (separate file)**
$f_4$-ratio statistics for tested scenarios.

**Table S4. (separate file)**
Admixture graph model fit statistics.

**Table S5. (separate file)**
Enriched annotation terms for genes identified via $f_{dM}$ from wild introgression into Zebu (*Bos t. indicus*).

**Table S6. (separate file)**
Enriched annotation terms for genes identified via $f_{dM}$ from domesticate introgression into Zebu (*Bos t. indicus*).

**Table S7. (separate file)**
Genes containing nonsynonymous mutations that matched the pattern of introgression into Zebu (*Bos t. indicus*).

**Table S8. (separate file)**
Enriched annotation terms for genes identified via $f_{dM}$ from wild introgression across three admixture events.

**Table S9. (separate file)**
Enriched annotation terms for genes identified via $f_{dM}$ from domesticate introgression across three admixture events.
